# Metabarcoding of sedimentary ancient DNA reveals centuries of species turnover and introductions of non-indigenous species in a highly urbanised estuary

**DOI:** 10.64898/2025.12.02.691833

**Authors:** Elena Baños, Luke E. Holman, Nicola Pratt, Andrew B. Cundy, Sandra Nogué, Marc Rius

**Affiliations:** Department of Marine Ecology, Centre for Advanced Studies of Blanes (CEAB), Spanish Research Council (CSIC), 17300 Blanes, Catalonia, Spain; Universitat de Barcelona, 08028 Barcelona, Catalonia, Spain; Section for Molecular Ecology and Evolution, Globe Institute, University of Copenhagen, Copenhagen, Denmark; School of Ocean and Earth Science, National Oceanography Centre Southampton, University of Southampton, Southampton, United Kingdom; GAU-Radioanalytical, School of Ocean and Earth Science, National Oceanography Centre Southampton, University of Southampton, Southampton, United Kingdom; Universitat Autònoma de Barcelona, Bellaterra (Cerdanyola del Vallès), 08193, Catalonia, Spain; CREAF, Bellaterra (Cerdanyola del Vallès), 08193, Catalonia, Spain; Department of Zoology, Centre for Ecological Genomics and Wildlife Conservation, University of Johannesburg, Auckland Park Johannesburg 2006, South Africa

**Keywords:** Biological invasions, early introductions, historical population dynamics, ancient environmental DNA

## Abstract

Understanding how anthropogenic pressures shape ecosystems over time is crucial for managing both biodiversity and bioinvasion risk. Despite recent research efforts focussing on temporal community changes as a result of human impacts, little is known about such changes over periods exceeding a century. Here, we reconstructed long-term community changes and non-indigenous species (NIS) dynamics by using sedimentary ancient DNA (sedaDNA) metabarcoding and geochemical analysis of sediment cores samples collected from a temperate estuary (Southampton Water, southern British Isles) subject to significant human impacts for centuries. We detected 73 species of Metazoa and Plantae and classified them by their status (native, cryptogenic and NIS). Our results revealed sporadic detection of NIS (both failed and continuously successful NIS introductions) throughout the entire sedimentary record. Notably, some NIS were detected as early as the 18^th^ century, indicating the presence of NIS introductions in pre-industrial times. We found a continuous overall decline in native species richness since the 19^th^ century. In addition, we found distinct peaks in copper and zinc concentrations reflecting recent periods of intensified industrial and port activity, highlighting the role of human activities in shaping biodiversity patterns and NIS dynamics. Our findings demonstrate how metabarcoding of sedaDNA archives can reveal trends in the distribution of NIS and wider biodiversity over centuries. The integrated approach used here provides new insights into biodiversity changes through time, unravels key information for biodiversity conservation management and biosecurity policy, and shows the potential that sedaDNA metabarcoding has for investigating intrinsic attributes of NIS introductions.

## Introduction

Understanding fine-scale biodiversity change through time is crucial in ecology and evolutionary biology, yet long-term records that capture pre-industrial baselines remain rare (Holman et al., 2025a). This gap limits our ability to distinguish natural variability from human-driven change, despite the accelerating influence of anthropogenic activities across ecosystems worldwide (Dornelas et al., 2014; Magurran et al., 2018; Nogué et al., 2021; Pereira et al., 2010; Wauchope et al., 2021). A full understanding of these long-term dynamics requires multidisciplinary approaches that combine complementary methods such as historical molecular data with geochemical and ecological analyses. Such multidisciplinary approach has unravelled the long-term effects of human activities, reconstructed biodiversity patterns over centuries (Garcés-Pastor et al., 2022; Giguet-Covex et al., 2014; Holman et al., 2025a), and detected non-indigenous species (NIS) introductions (Ficetola et al., 2018; van Leeuwen et al., 2008; Walentowitz et al., 2023). However, no study to date has successfully integrated molecular palaeoecology with invasion science to study a large number of NIS in highly urbanised ecosystems.

Invasive species are among the greatest threats to global biodiversity, exerting profound ecological and socioeconomic impacts across terrestrial and aquatic ecosystems (Simberloff et al., 2013). They can disrupt food webs, outcompete native taxa, and impact fisheries and aquaculture operations (Turbelin et al., 2022). These introductions set in motion cascading impacts at both ecological and societal levels (Vilà & Hulme, 2017). The spread of NIS is tightly linked to the frequency and intensity of anthropogenic activities such as maritime transport and port construction (Bishop et al., 2017; Darling & Carlton, 2018; Hudson et al., 2022), and result in novel environmental changes and biotic interactions (Ficetola et al., 2018; Johnston et al., 2017). Most studies focus on contemporary monitoring of NIS (Haubrock & Soto, 2023), leaving both historical NIS dynamics and community-level effects poorly understood (Miura, 2007). This is somewhat surprising, as the temporal context is essential to understand range expansions, niche shifts, and community changes (Gallardo et al., 2016; Rius et al., 2015; Sorte et al., 2010). Understanding when, how, and under what conditions NIS arrive and persist is crucial, yet remains a relatively underexplored aspect of invasion science (Catford et al., 2009; Ricciardi et al., 2017). Greater attention to this topic could enhance future predictions of NIS spread and inform strategies to mitigate their community impacts.

Molecular tools such as metabarcoding and metagenomics have in recent years revolutionised the detection and monitoring of biodiversity (Holman et al., 2025b), including NIS (Bailey et al., 2020; Zarcero et al., 2024). The study of environmental DNA (eDNA), defined as genetic material shed into the environment from cells, tissues, gametes, or carcasses, can be recovered through non-invasive sampling approaches (e.g., water filtration, sediment sampling) and subsequently analysed to detect species across taxonomic groups, including metazoans (Ficetola et al., 2008), fungi, and protists (Cowart et al., 2018; Mahon et al., 2013). eDNA persistence varies widely, from hours in water (Collins et al., 2018; Holman et al., 2022) to hundreds of thousands or even millions of years in sediments and soils (Kjær et al., 2022; Parducci et al., 2017; Sakata et al., 2020), providing temporal depth to biodiversity assessments (Stager et al., 2015). One type of eDNA is sedimentary ancient DNA (sedaDNA), which is increasingly recognised as a powerful tool for reconstructing past ecological communities and environmental change (Capo et al., 2021; Holman et al., 2025a; Nguyen et al., 2023). Marine intertidal zones, in particular, offer ideal conditions for sedaDNA preservation due to relatively stable sediment accumulation, periodic sub-oxia or anoxia, and high organic content (Alsos et al., 2023; Cundy et al., 1997). These settings are also among the most impacted by anthropogenic pressures, including ballast water discharge, port operations, and coastal development (Bishop et al., 2015; Guillemaud et al., 2011), making them strategic natural archives for investigating long-term biological invasions. When placed within a chronological framework derived from sedimentary dating and geochemical profiles (e.g. radiocarbon, ^210^Pb, ^137^Cs) (Appleby, 2002; Goslar et al., 2005), sedaDNA records provide unprecedented opportunities to trace community change and ecological disturbances over centennial to millennial timescales (Capo et al., 2021; Parducci et al., 2017). This chronological resolution makes sedaDNA a particularly valuable tool for detecting both long-term biodiversity trends and the timing of ecological disturbances.

The study of sedaDNA in biodiversity research and invasion science remains limited (Ficetola et al., 2018; Miura, 2007; Pérez et al., 2023). While some historical trends in marine bioinvasions have been explored (Carlton & Schwindt, 2024), very few studies have provided spatio-temporally constrained reconstructions of NIS arrivals, pinpointing when and where specific taxa first appeared. Here, sedaDNA offers a unique opportunity to recover what was thought lost: direct molecular evidence of species introductions at defined points in time and space. Recent advances in high-throughput metabarcoding and palaeogenomic approaches (Holman et al., 2025b) now allow us to address key questions such as: When did NIS first arrive? What is the impact of priority effects on NIS arrival? How did NIS presence correlate with environmental change? What role have human activities played in shaping contemporaneous communities?

Here, we used sedaDNA metabarcoding and geochemical analysis to study sediment cores from a historically industrialised and ecologically significant marine intertidal ecosystem. We integrate data from different genetic markers with analyses of trace metals to:

1. Reconstruct temporal patterns in species richness and community composition.
2. Detect native, cryptogenic and NIS across the sediment record.
3. Identify potential links between anthropogenic activities (here, indicated by metal pollution during industrialisation) and community change.
4. Assess the value of intertidal sedaDNA archives for long-term bioinvasions monitoring.

We hypothesised that increased maritime activity during the mid-20^th^ century has contributed to a rise in NIS richness and has significantly altered community composition. In addition, we expected trace metal (e.g. Cu, Zn and Pb) concentrations to reflect industrial and shipping pressures that may correlate with community shifts. By bridging molecular palaeoecology and invasion science, we expected to unravel the temporal dynamics of NIS (e.g. detection of failed introductions) and intrinsic characteristics of the ongoing transformation of estuarine ecosystems under human influence.

## Methods

### Study region

Our study focused on two intertidal zones within the wider Southampton Water estuarine system, United Kingdom, the first located at Hythe in the main body of Southampton Water, and the second at Bursledon (Fig. 1), 5.5 km upstream from the mouth of the Hamble estuary (see Supplementary Table 1). The Southampton Water is a mesotidal, temperate, coastal plain estuary (covering an area of approximately 20 km^2^) which receives freshwater discharges from the Test, Itchen, Hamble, and Meon Rivers (Celis-Hernandez et al., 2022). Despite these freshwater inputs, the Southampton Water is a tidally dominated, well-mixed system, with salinities typically ranging from 25 to 35 PSU and only limited vertical stratification under normal conditions (Levasseur et al., 2007). The estuary exhibits a pronounced gradient in both hydrodynamics and sediment characteristics, with stronger tidal currents and coarser sediments in the lower estuary near Hythe, and finer, more organic-rich muds deposited in the upper reaches of the Hamble estuary near Bursledon (Guo & Zhang, 2010).

**Fig. 1.**
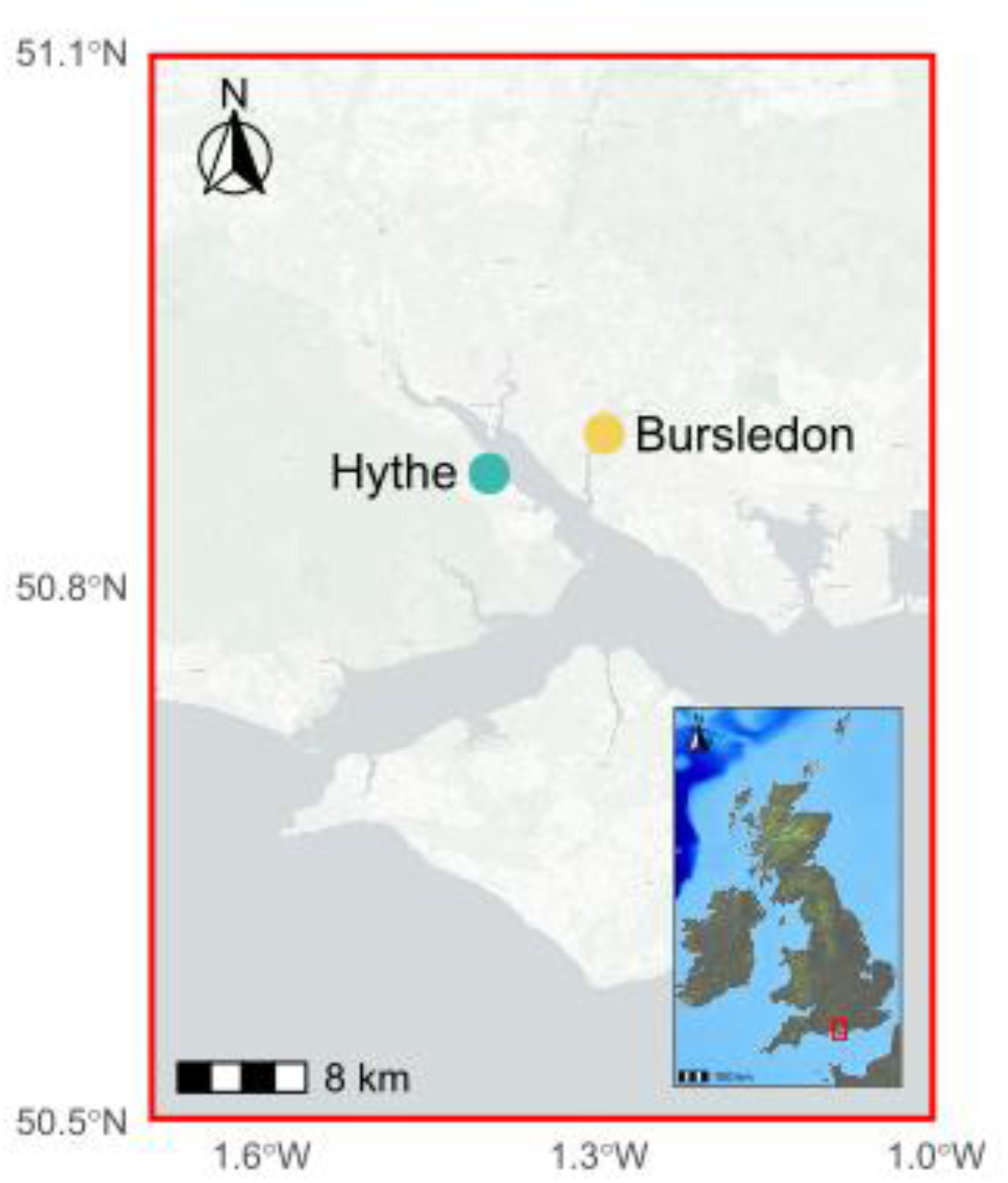
Location of Southampton Water within the southern British Isles (inset), and locations of the sampling sites at Hythe (blue circle) and Bursledon (yellow circle, in the Hamble river). In order to illustrate the geographic setting of Southampton Water and the sampling sites, we used *marmap* package (v1.1.10) (Pante & Simon-Bouhet, 2013).

The wider Southampton Water estuary has a long history of port, urban and industrial development (Celis-Hernandez et al., 2022) and has previously been studied to investigate the retention of trace elements and isotopic Pb sedimentary signatures in a system experiencing significant sea level rise, yielding reliable geological sediment records (Cundy et al., 1997, 2003; Cundy & Croudace, 2017).

### Core sampling and subsampling

Two sediment cores (HM01 and HB5) with a diameter of 50 mm (of 30cm (HB5) and 50cm (HM01) length) were collected using a peat borer (UWITEC, Mondsee, Austria), while a longer (90cm length) core was physically cut out of the marsh face as a monolith (HM03). The cores were collected at the following sampling sites: HM01 (UTM: 619883.1, 5638889.9) and HM03 (UTM: 613427.4, 5635923.5) at Hythe, and HB5 (UTM: 613499.9, 5635915.9) at Bursledon (Supplementary Table 1). In order to avoid contamination, all tools, including PVC tubes, scissors and the peat borer were sterilised using 10% sodium hypochlorite prior to sample collection. The cores were wrapped with plastic film upon collection and transported to a controlled temperature room at 4 °C at the School of Geography and Environmental Science (SOGES), University of Southampton.

Subsampling was conducted in a dedicated PCR-free laboratory at SOGES. The outermost millimetres of each core were removed using sterile disposable scalpels and spatulas to avoid contamination risk. Sediment subsamples were collected using sterile disposable scalpels and spatulas and were transferred to sterile 50 mL Falcon tubes and stored at −20°C. Disposable gloves were changed between each subsample collection, and reusable equipment was decontaminated using first 10% bleach and then 70% ethanol. In total, 56 subsamples were obtained: 35 from HM01 (sampled at 1 cm intervals), 6 from HM03 (every 20 cm), and 15 from HB5 (every 2 cm). Sampling intervals varied among cores to account for differences in sediment accumulation rates, total core length, and material availability, aiming to achieve approximately comparable temporal resolution across sites while ensuring sufficient sediment for sedaDNA, dating, and geochemical and radiometric analyses (Nguyen et al., 2023).

### Dating and X-Ray fluorescence core scanner

XRF-CS analysis was used to determine the concentration of trace elements throughout the two sedimentary records. Hythe cores were analysed using ITRAX X-ray fluorescence core scanning (XRF-CS) at the British Ocean Sediment Core Research Facility (BOSCORF, National Oceanography Centre, Southampton). Each half core was scanned using an Itrax micro-XRF scanner (step size: 200 μm; counting time: 30 s; Mo anode X-ray tube at 30 kV, 50 mA), which produced intensity data (counts) for 54 trace elements in total, following Cundy & Croudace, 2017. This non-destructive method allows for high-resolution elemental analysis of sediment cores (Croudace et al., 2019). The elemental concentrations were recorded in counts per second (CPS), data were normalized to total scatter to account for changes in water content and sediment density following Kylander et al., 2011 and Fisher et al., 2025) (Supplementary Data S1). The XRF data from the Bursledon site (core HB5) was extracted from Appelt et al. (submitted) using the same methodology as above.

Sediment chronology was determined through radiometric dating of 42 samples, comprising 15 from HM01, 14 from HM03, and 13 from HB5, via measurement of ^137^Cs and ^210^Pb. Radionuclide activities were determined in freeze-dried, lightly ground samples via γ-ray spectrometry using HPGe well-type detectors (Mirion Technologies, Hampshire, UK). ^210^Pb derived ages were calculated following the Constant Flux-Constant Sedimentation Rate (or Simple) model (Appleby, 2002; Appleby & Oldfield, 1978; Croudace & Cundy, 1995), while ^137^Cs-derived ages were calculated using the position of the prominent 1963 fallout maximum from nuclear weapons testing, following Cundy et al., 1997 and Cundy & Croudace 2017 (Fig. S2).

### DNA extraction

We extracted sedaDNA from the intertidal core samples at a dedicated PCR-free laboratory at the SOGES. A total of 41 samples were processed (15 from HM01, 5 from HM03, 13 from HB5, and 8 negative controls). Prior to use, all surfaces in the PCR-free laboratory at SOGES were cleaned with 10% commercial bleach solution (approximately 0.5–0.6% sodium hypochlorite), followed by 70% ethanol to avoid contamination by high-copy DNA templates.

DNA was extracted using the DNeasy PowerMax Soil Kit (QIAGEN, Netherlands), following the manufacturer’s protocol with the following amendments to Steps 1-4 and 19. PowerMax Bead Tubes containing 15 ml of PowerBead solution and 10 g of sediment subsample were vortexed at max speed for 2 minutes, 1.2 ml of Solution C1 was added, tubes were vortexed for 30 seconds and then placed in a rotating hybridisation oven for 30 mins at 65°C, 35 rpm. DNA was eluted into 3 ml Solution C6. Negative controls were included at every step of the workflow, including DNA extraction (n=6) and PCR amplification (n=8 per library) a total of 30 negative controls. Extracted DNA was quantified using the Qubit HS Assay Kit (Thermo Fisher Scientific) and 80 µL were purified with the OneStep PCR Inhibitor Removal Kit (ZYMO Research, USA) to ensure the effective removal of potential contaminants that could impede downstream enzymatic reactions. DNA extractions were stored at −20°C until they were processed.

### Primer selection and library preparation for metabarcoding samples

Two primer sets were used for metabarcoding: The first targeted a 313 bp fragment of the cytochrome c oxidase I (COI) gene using the Leray-XT primers (Wangensteen et al., 2018), which consisted of mlCOIintF-XT (5′-GGWACWRGWTGRACWITITAYCCYCC-3′) and jgHCO2198 (5′-TAIACYTCIGGRTGICCRAARAAYCA-3′), while the second primer set targeted the V7 region of the 18S rRNA gene (∼110 bp) using the 18S_allshort primers (Guardiola et al., 2015), with a forward 5′-TTTGTCTGSTTAATTSCG-3′ and a reverse 5′-TCACAGACCTGTTATTGC-3′. All amplicons were dual-indexed using a twin-tagging scheme, with 8-bp tags (minimum Hamming distance 3 between tag barcodes). appended to the 5′ end of both forward and reverse primers. For each sample, the same tag sequence was used on both primers (i.e. mirrored tags) to reduce tag-jump misassignment (Schnell et al., 2015).

For each gene, we amplified and sequenced eight independent technical replicates of 40 µL reactions, with 4 µL of sedaDNA per replicate (32 µL total per sample). Each PCR mix contained: 20 µL AmpliTaq Gold 360 Master Mix, 11.6 µL bovine serum albumin (BSA; 20 mg/mL, Fisher Scientific), 2 µL of each primer (5 nmol/mL), and 4 µL of sedaDNA extract. PCR conditions for COI were 95°C for 10 min; 35 cycles of 94°C (1 min), 45°C (1 min), 72°C (1 min); final extension at 72°C (5 min). For 18S, we used 95°C for 10 min; 35 cycles of 95°C (30 s), 45°C (30 s), 72°C (30 s); final extension at 72°C (5 min). Amplicons were purified using the MinElute PCR Purification Kit (QIAGEN). Samples were quantified using Qubit and pooled at equimolar concentrations. A total of four libraries were prepared for each marker. Each sequencing pool comprised 96 samples, including 8 negative controls, and libraries were prepared using the TruSeq PCR-Free Library Prep Kit (Illumina) was used with a 0.13 bead ratio (COI) and 0.16 (18S). Illumina P5 and P7 adapters were used for library preparation.

Sequencing was performed on an Illumina NovaSeq 6000 platform using 2 × 250 bp paired-end reads for COI (SP flow cell) and 2 × 150 bp paired-end reads for 18S (S4 flow cell), targeting approximately one million reads per sample.

### Bioinformatics

Raw paired-end reads were demultiplexed by sample-specific dual barcodes using Cutadapt v2.3 (Martin, 2011). Barcodes for the forward and reverse reads were supplied as tag files for sense and antisense orientations, and demultiplexing was performed with --pair-adapters and --pair-filter=both, such that both mates in a pair were required to carry the correct barcode combination. We allowed up to 10% mismatches in barcode sequences (--error-rate 0.1) while disallowing insertions and deletions (--no-indels), and reads lacking a valid barcode pair were written to separate “unassigned” files. DADA2 v1.12 (Callahan et al., 2016) was used for the following denoising and cleaning steps. Reads with >1 expected error were discarded. Trimming was based on quality: forward reads were trimmed to 250 bp (COI) and 120 bp (18S); reverse reads to 230 bp (COI) and 110 bp (18S). Merged reads were retained only if within the expected fragment length (303–323 bp COI; 100–200 bp 18S). Subsequently, chimaeras were removed. Amplicon sequence variants (ASVs) were curated using the LULU algorithm (Frøslev et al., 2017) in the metabarTOAD pipeline (available at https://github.com/leholman/metabarTOAD/blob/master/R/ApplyLulu.R), which employs *vsearch* for pairwise similarity searches (Rognes et al., 2016), using parameters of a minimum match threshold of 98%, a *vsearch* identity cutoff of 0.84, a minimum query coverage of 0.9, and a maximum of 10 hits per query.

COI and 18S ASVs were taxonomically assigned using BLAST+ v2.14.1 (Camacho et al., 2009) against the NCBI nt database (downloaded 27^th^ of March 2024), sequences were then parsed using *ParseTaxonomy* function in R (metabarTOAD), which (i) removes hits labelled as uncultured/environmental/clone and epithets “sp.”; (ii) retains high-confidence assignments above user-defined thresholds of percent identity and coverage; (iii) when multiple high-confidence taxIDs exist for an ASV, resolves the rank via a lowest common ancestor (LCA) using the NCBI lineage file; and (iv) otherwise records low-confidence hits above relaxed thresholds as best-bitscore matches. For COI, we set high-confidence thresholds to ≥97% identity and ≥85% coverage (both query and HSP), and low-confidence thresholds to ≥85% identity and ≥65% coverage. For 18S, species-level assignments were accepted only at 100% identity (≥98% coverage); when unmet, genus-level assignments were accepted at ≥99% identity (≥98% coverage) using the same parsing/LCA logic. Ambiguous or environmental hits were discarded.

Quality-control filtering of the ASV-by-sample table generated by DADA2 involved several steps to ensure robust and reliable downstream analysis. Firstly, a minimum threshold of three reads per observation was applied to eliminate low-frequency artefacts. ASVs detected only once across all replicates were discarded. To address potential contamination, the maximum read count observed among all extraction and PCR negative controls was used as a baseline threshold, whereby any read counts in non-control samples that fell below this value were set to zero. Following this, both COI and 18S datasets were filtered to retain only taxonomically informative ASVs, defined as those (i) meeting the sequence identity and coverage thresholds established in the taxonomic assignment step, (ii) classified to at least phylum level, and (iii) not labelled as “uncultured”, “environmental sample”, or “metagenome”. Remaining ASVs were then subset by eukaryotic group: for COI, metazoans were retained; for 18S, metazoans, plants, and protists were retained, while bacterial and unclassified sequences were excluded. These curated datasets were used for conducting separate or combined analyses according to the relevant taxonomic groupings. Within each dataset, the read counts were converted to relative abundance by technical replicates, and these replicates were then collapsed to yield the mean relative abundance per ASV per sample. The processed sedaDNA datasets, consisting of total and relative abundance matrices of filtered ASVs for COI and 18S genes across all samples, are provided in Supplementary Data S2.1 and S2.2 for total abundances and Supplementary Data 2 for relative abundances.

All metazoan and plantae species identified in the dataset were manually classified into biogeographic status based on a comprehensive literature review (detailed in Supplementary Data S3). Species were designated as NIS if their origin was known and were allochthonous to the Solent Estuary, as cryptogenic species (CRY) if their origin could not be determined but their introduced status in the study region was certain, as native species (NAT) when their presence was confirmed to be endemic, and as species of unknown status (UNK) if the origin or the introduced status could not be determined.

### Statistical analysis and data interpretation

All statistical analyses and data visualisations were performed in R (v4.3.1). Species richness, community structure, and taxonomic composition were evaluated using functions from the *vegan* package (Oksanen et al., 2009). Species richness was calculated based on the total number of unique ASVs per sample after quality filtering, without rarefaction, and compared across sites and sediment layers to assess spatial and vertical diversity patterns.

To explore differences in community composition, non-metric multidimensional scaling (nMDS) was carried out using Bray–Curtis dissimilarity matrices. The ordination results were used to assess clustering by core location, depth, and marker type (COI and 18S). Taxonomic composition was visualised using barplots displaying relative abundance of major taxonomic groups (e.g., phyla or classes) across samples. These visual summaries helped identify key differences in assemblages between sampling sites and core depth intervals.

In addition, we used general linear models (ordinary least squares) in R using the *lm* function from the *stats* package (v.3.6.2). Model structure followed the form Richness ∼ Site × Marker, where Site (Hythe, Bursledon) and Marker (COI, 18S) were treated as categorical explanatory variables. To investigate temporal patterns in species richness across status categories, we applied Generalised Additive Models (GAMs) using the gam function from the *mgcv* package (v1.9-0) to visualise the historical trajectories of NIS, NAT, CRY and UNK at each site (Fig. 4).

All figures and plots were generated using *ggplot2* (Wickham, 2011), and compositional data were handled with the aid of the *phyloseq* package (McMurdie & Holmes, 2013) for downstream visualisation and filtering.

## Results

### Sample dating

The ^210^Pb profile in core HM01 shows a shallow subsurface activity maximum followed by a broadly exponential decline with depth (Fig.S1). Application of the Constant Flux–Constant Sedimentation Rate (Simple) model (Appleby, 2002) indicates a sediment accumulation rate (SAR) of 4.8 mm y⁻¹. The prominent subsurface ^137^ Cs activity maximum at approximately −27 cm depth, attributed to the 1963 nuclear weapons testing fallout maximum (Cundy & Croudace, 2017), yields a SAR of 4.9 mm y⁻¹, in excellent agreement with the ^210^Pb-derived rate. Core HM03, a longer monolith collected from the unvegetated marsh edge, showed physical evidence of recent surface erosion (e.g., vegetation removal and wave scouring). ^210^Pb activities remained close to supported levels (estimated from ^214^Pb) of ∼20 Bq kg⁻¹ throughout the core, and ^137^Cs was only detected in the surface sample, indicating the loss of surface sediment and the very recent depositional record (Cundy et al., 2003). Dates older than 1900 were estimated by extrapolating the SARs derived from radiometric and geochemical profiles, assuming approximately uniform sedimentation rates over time.

In core HB5 (Bursledon), ^210^Pb shows a near-uniform decline with depth, consistent with a SAR of 4.1 mm y⁻¹ (derived from the Simple model). The ^137^ Cs activity profile displays a broad maximum between −20 and −27 cm depth, interpreted as the 1963 nuclear weapons fallout peak, with possible contributions from earlier events (e.g., 1958; Appelt et al., submitted).

### XRF core scanner

XRF data for contaminant elements with well-defined temporal input trends into Southampton Water, particularly copper (Cu) and lead (Pb) (Cundy & Croudace, 2017), provided additional chronological control, especially for the Hythe cores HM01 and HM03 (Fig. S2). Accumulating saltmarsh sediments around Southampton Water retain a prominent Cu maximum linked to discharges from the Exxon oil refinery at Fawley, dated to 1970–1971 (Cundy et al., 1997; Cundy & Croudace, 2017). Pb similarly exhibits a broad maximum associated with increased urban and industrial activity around 1940 (Cundy & Croudace, 2017).

In core HM01, the Cu maximum at −26 cm corresponds to 1970–1971 and implies a SAR of 5.4 mm y⁻¹, in close agreement with the radiometric results (Fig.S2). Zn displays a broad maximum between −10 and −20 cm, possibly reflecting more recent industrial or shipping inputs; however, Zn mobilisation by early-diagenetic processes at Hythe may complicate its use as a chronological tracer (Cundy & Croudace, 2017). The Pb profile shows two broad maxima, at −35 cm and −20 cm, likely corresponding to the 1940 Pb enrichment and increased Pb inputs from gasoline combustion during the 1970s, respectively.

Despite surface erosion, core HM03 retains part of the 1960s Cu increase associated with Fawley refinery expansion and the 1940 Pb maximum, the latter coinciding with a Zn enrichment (as also noted by Cundy & Croudace, 2017). The Pb maximum occurs at −14 cm, suggesting 14 cm of sediment accumulation in ∼27 years, equivalent to ∼5.2 mm y⁻¹, consistent with the rate observed in the undisturbed HM01 core.

In contrast, in the Bursledon core (HB5), localised inputs (including runoff from the motorway bridge opened in 1976, particularly for Pb) have obscured these broader regional geochemical trends (Appelt et al., submitted).

### Community structure detection from sedaDNA

Following rigorous filtering and quality control of the metabarcoding datasets, a total of 7,199 ASVs were retained for the COI marker, of which 794 could be taxonomically assigned. Among retained ASVs, 83.4% were attributed to eukaryotes and 16.6% to bacterial sequences. For the 18S marker, 36,321 ASVs were retained, with 3,949 successfully assigned to a taxonomic lineage. Within this subset, 99.95% were identified as eukaryotic, and 0.05% corresponded to bacterial taxa.

Analysis of genus-level richness across sediment layers revealed consistent detection of diverse taxa over time, with no observable decline in richness in older samples (Fig. 2, centre panels). In fact, certain historical layers (e.g., 1950 and 1965 at Bursledon) exhibited richness levels comparable to or higher than recent samples. These findings suggest strong DNA preservation and stable detection power across the sedimentary record for both COI and 18S datasets. COI generally showed lower richness values than 18S (Fig. 2A), but similar trends were evident between markers, especially in the Bursledon core (Fig. 2A).

**Fig. 2.**
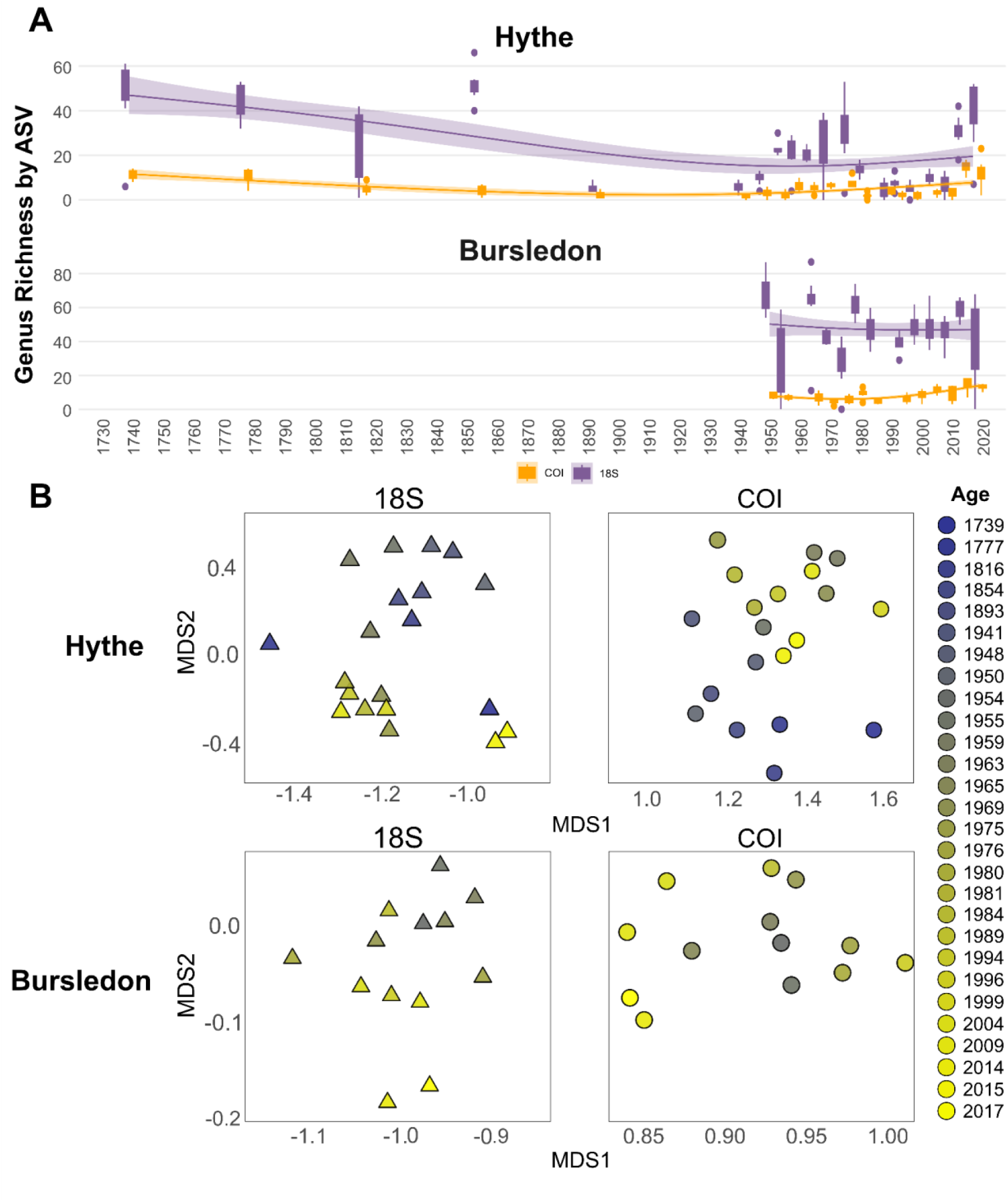
(A) Temporal variation of genus richness at each sampling site (i.e. Hythe and Bursledon) considering the datasets of amplicon sequence variants (ASVs) collapsed by genus. These datasets were obtained using two genetic markers: a section of the cytochrome c oxidase I (COI) gene and a section of the 18S rRNA gene (18S). Boxplots represent richness values by dated sediment layers; lines show smoothed richness trends through time. (B) Non-metric multidimensional scaling (nMDS) plots based on Bray–Curtis dissimilarity matrices for each sampling site, illustrating differences in taxonomic community structure between cores and marker types (COI: circles; 18S: triangles).

Community composition was further explored through nMDS based on Bray–Curtis dissimilarities. The ordination plots (Fig. 2B) revealed clear separation between the two genetic markers (COI and 18S) and among sediment layers within each core. Samples clustered by gene, and the second nMDS axis (MDS2 in Fig. 2B) was strongly associated with depth/age, suggesting temporal turnover in community composition at both sampling sites.

Our GLM analysis confirmed that richness differed significantly between cores for both genetic markers (Table S2a). The model also revealed significant effects of sampling year and environment (estuary vs river), although the interaction term was not always significant, significant differences emerge mainly in recent decades (Table S2b).

Our broad taxonomic analysis of the relative abundance of reads at the kingdom level for both the COI and 18S datasets showed marked temporal and spatial shifts in community composition across the study sites (Fig. 3). For the COI dataset, Protists were the most abundant kingdom across nearly all sediment layers and both sites, with particularly high representation in mid-20^th^ century samples. Plantae (mostly Viridiplantae) and Fungi were consistently detected across the record, although less consistently than in the 18S dataset. Metazoan reads were also present throughout. Samples from the more riverine core (Bursledon) generally exhibited higher proportions of Protist reads compared to the fully estuarine core (Hythe).

**Fig. 3.**
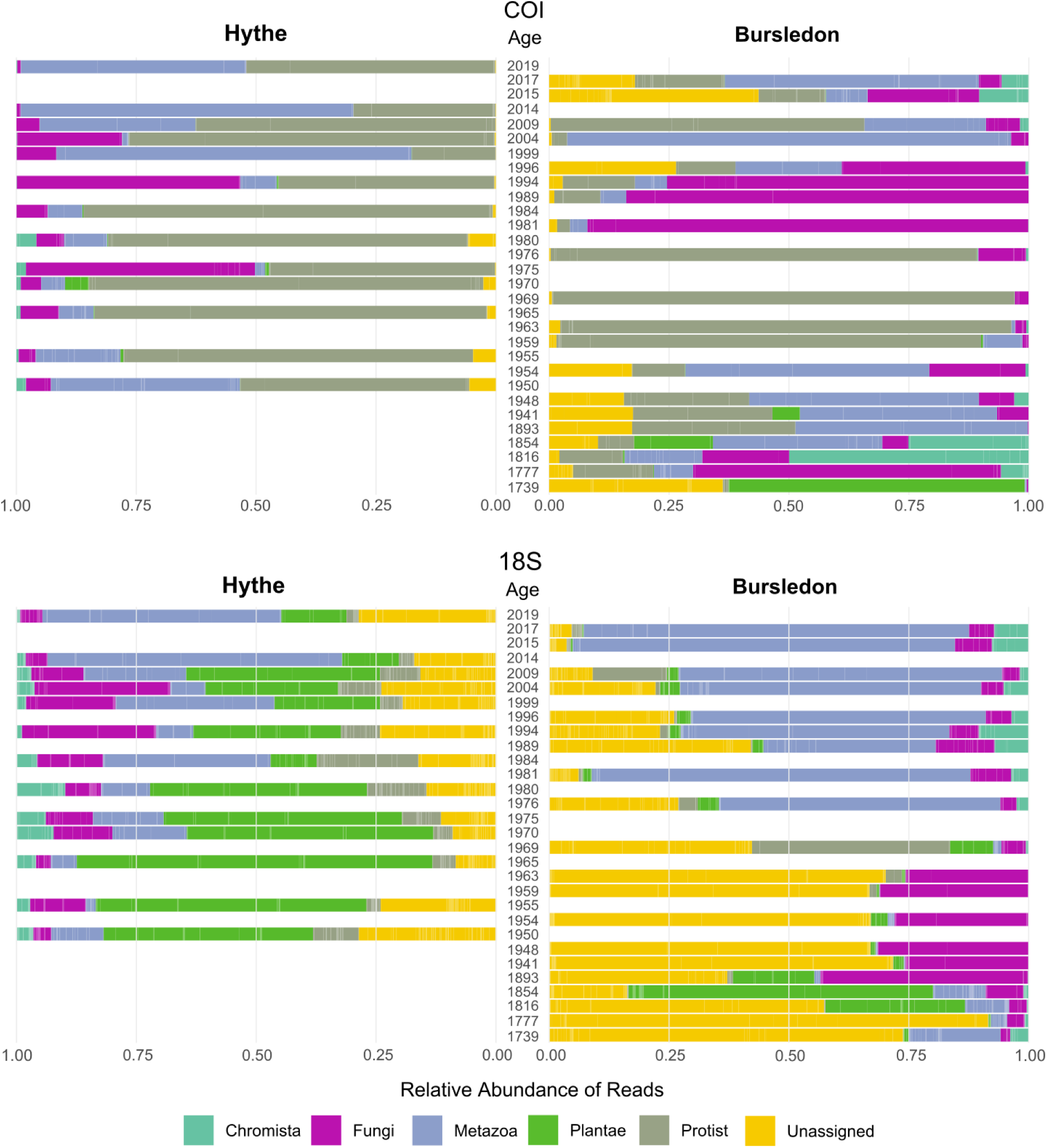
Relative read abundance (percentage of total reads per sample) by kingdom across dated sediment layers for the sections of the analysed genes [cytochrome c oxidase I (COI) and 18S rRNA gene (18S)] respectively. Samples are grouped by site (Hythe and Bursledon) and ordered chronologically. The six kingdoms with the highest mean relative read abundance across all samples are shown, including Protist, Plantae, Fungi, Chromista, Metazoa, and Unassigned sequences.

The 18S dataset displayed complementary yet distinct patterns (Fig. 3). Plantae reads dominated most of the Bursledon samples, particularly in the oldest layers, whereas Fungal reads were more prevalent in the Hythe site. Chromista sequences showed occasional peaks, and Metazoan reads, though consistently present, appeared in lower relative abundance compared to the COI dataset. The relatively lower proportion of Animalia in 18S likely reflects primer bias or differences in amplification efficiency. Nonetheless, consistent temporal and spatial patterns across both markers suggest similar underlying ecological trends.

Our exploration of faunal dynamics examining metazoan reads at a phylum level across time using both markers showed a dominance of Annelida, Nematoda, and Platyhelminthes, particularly in shallow and more recent layers (Fig. S3). Although these groups were common in soft-sediment habitats, their abundance and consistency across depths suggested a long-term faunal continuity in the system. Arthropoda and Mollusca were also detected but in more restricted intervals, possibly reflecting habitat changes or differential preservation. Notably, Hythe exhibited a broader representation of phyla over time compared to the Bursledon core, hinting at more heterogeneous or temporally dynamic benthic conditions in the study sites.

All the 73 identified species belonged to the kingdom Metazoa and Plantae (Supplementary Data S4). A phylum-by-kingdom breakdown of these classified species is provided in Fig. S4.

GAMs showed that the smooth term for unknown classified species richness was significantly different from a flat line at both sites (Fig. 4; Fig. S5; Table S3). At Hythe, the smooth function for NAT richness also showed a significant deviation from a flat temporal trend. In contrast, for NIS and CRY, the estimated smooth functions were not significantly different from a flat line, suggesting no clear temporal structure in richness for these categories.

**Fig. 4.**
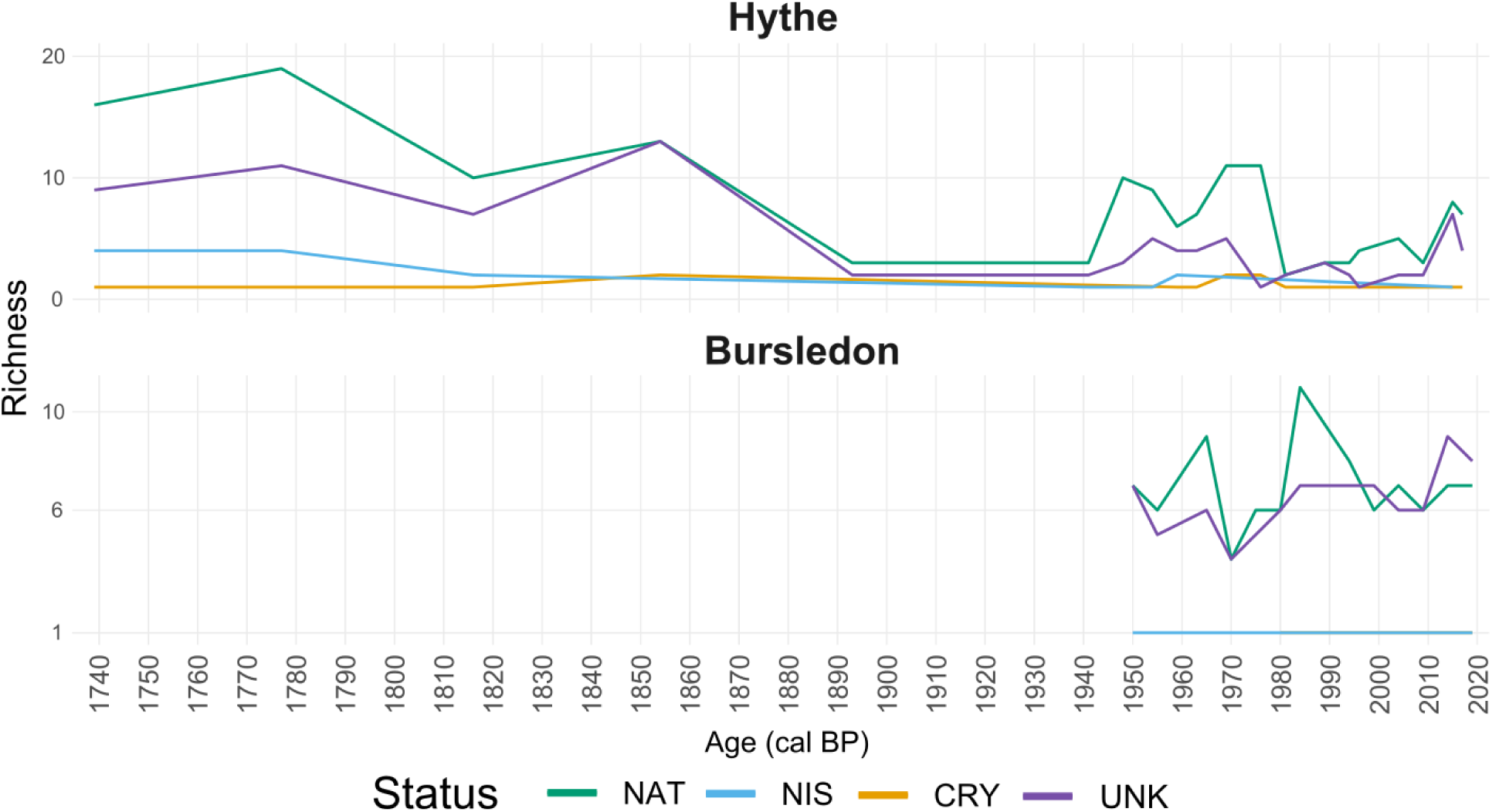
Temporal trends in species richness by biogeographic status in Hythe and Bursledon cores. Representing the richness over time of non-indigenous species (NIS), native species (NAT), cryptogenic species (CRY), and taxa with unknown status (UNK). Note that the x-axis in the Bursledon panel starts at 1950 due to the sediment record range in that core.

At Hythe, native species richness exhibited a general decline from the early 18^th^ century to the present (Fig. 4). NIS also showed a decreasing trend over the same period, while cryptogenic species remained consistently low and stable. The richness of unknown followed a trajectory similar to that of native species for most of the record but diverged in the most recent decades, suggesting a possible shift in community composition or detection bias. In contrast, the Bursledon core revealed a different pattern: NAT and unknown both displayed substantial richness throughout the record, with their trajectories alternating in dominance over time. Meanwhile, the richness of NIS remained consistently low, with typically only a single observation per year. No CRY were detected at this site.

In order to examine the potential influence of historical human activity on NIS emergence, we compared the temporal dynamics of four focal NIS together with XRF-derived concentrations of the common contaminant metals copper (Cu), lead (Pb), and zinc (Zn) (Fig. 5; Table S4). The appearance of *Neoporphyra haitanensis* and *Nematostella vectensis* in the early 18^th^ century as an early NIS associated with maritime traffic, was detected only in the most recent sediment layers.

**Fig. 5.**
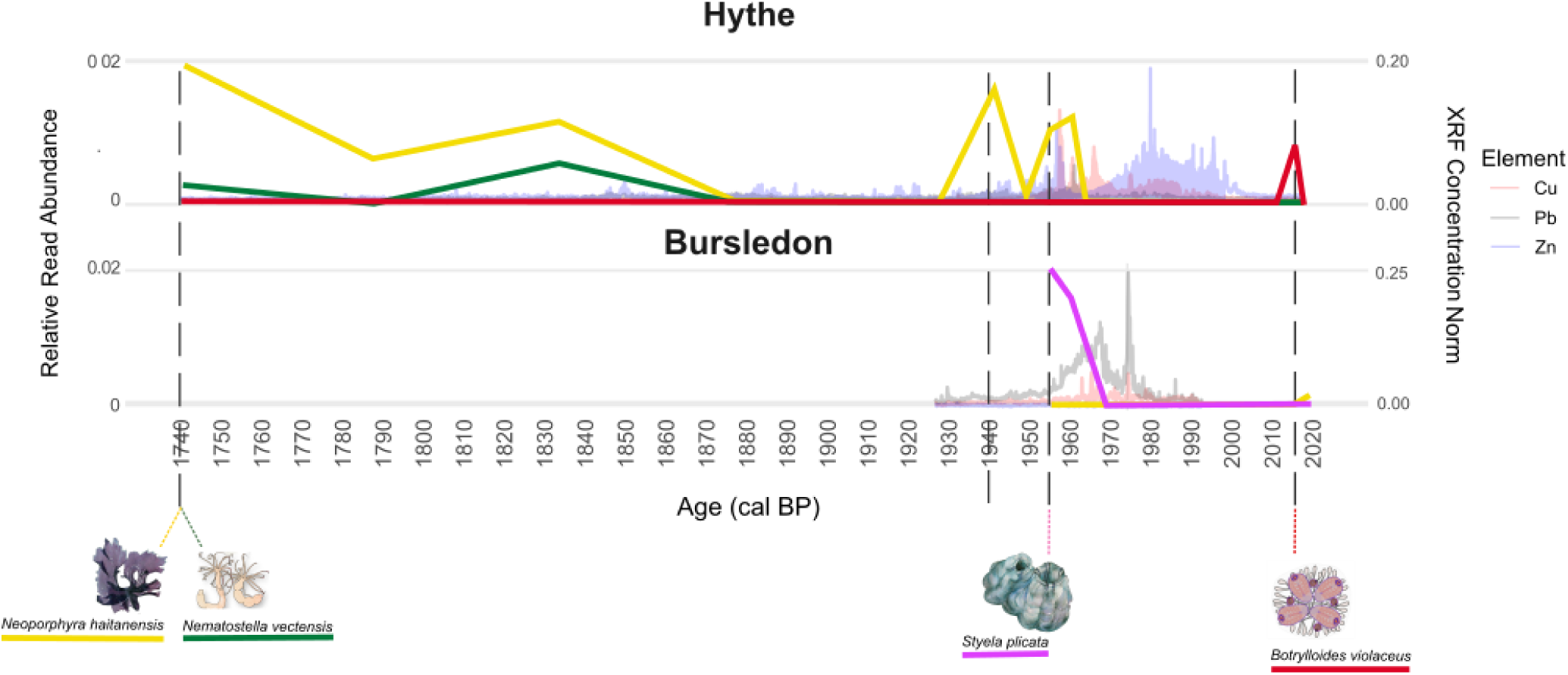
Relative read abundance of four focal non-indigenous species (*Neoporphyra haitanensis*, *Nematostella vectensis*, *Styela plicata* and *Botrylloides violaceus*) (Table S4) plotted against XRF-derived elemental concentrations of copper (Cu), lead (Pb), and zinc (Zn) from sediment cores. Both *N. haitanensis* and *N. vectensis* were detected using a genetic marker targeting a section of the 18S rRNA gene, while for *S. plicata* and *B. violaceus*, we amplified a section of the cytochrome c oxidase I gene.

Discrete maxima in Cu and Zn concentrations correlate historically with the major events of construction and marine traffic of the port of Southampton and temporally with the emergence or proliferation of several NIS species. The results for Pb showed a broader distribution but also peaked near industrial milestones (Fig. S2).

## Discussion

Our study revealed the temporal trends in composition of marine communities from intertidal areas spanning the past three centuries (*ca*. 1730 - 2020). Such long-term reconstructions are rarely achieved in marine environments due to the scarcity of studies assessing continuous historical records (Capo et al., 2021; Domaizon et al., 2017). We confidently detected Metazoa and Plantae species, with NIS being present from the 18^th^ century onwards (Fig. 5), indicating that estuarine introductions have occurred for centuries and that species translocations and turnover processes are long-standing (Carlton, 1996; Carlton & Schwindt, 2024; Reitzel et al., 2008). Regarding whole community richness patterns, native diversity declined through time since the 18^th^ century (Fig. S6), possibly linked to the increase in human activities following the Industrial Revolution, such as port construction, metal (and other) contamination, and maritime transport. This signal of industrialisation was evidenced by higher Cu and Pb concentrations observed in the XRF profiles (Fig. S2) (Albano et al., 2021; Leigh et al., 2019; Lotze et al., 2006; Turner, 2010). In addition, the riverine core (Bursledon) yielded a comparatively higher proportion of protist and plant reads than the fully estuarine core (Hythe) (Fig. 3), consistent with slight salinity differences between sites, as the Hamble receives greater freshwater input despite both sites being dominantly marine (Bodur et al., 2025; Jiang et al., 2022; Levasseur, 2008). Taken together, our results showed how the approach used here offers valuable insights into long-term biodiversity changes and dynamics of NIS introductions, providing new research opportunities for advancing knowledge in community ecology, biodiversity conservation, and invasion science.

### Reconstructing three centuries of marine biodiversity and invasion dynamics

In terms of historical biodiversity changes, sedaDNA preserved in intertidal and riverine mudflats retained a rich and structured record of biodiversity across several centuries, in line with observations from other depositional environments such as lakes, fjords, and marine basins (Alsos et al., 2023; Capo et al., 2021; Domaizon et al., 2017; Ficetola et al., 2018; Parducci et al., 2017; Pawłowska et al., 2014). Although sedaDNA is known to degrade with depth and age (Armbrecht et al., 2021; 2020), our results, extending across three centuries, showed no substantial richness and diversity shifts through time (Fig. 2) potentially linked to DNA degradation, suggesting that intertidal sediments can preserve sedaDNA effectively over centennial scales (Nguyen et al., 2023; Wang et al., 2021).

We found no significant increase in NIS richness over recent decades, contrary to common expectations of a continued rise driven by intensified maritime traffic, urbanisation, and port expansion (Bailey et al., 2020; Bishop et al., 2015; Sardain et al., 2019; Seebens et al., 2017). At the Hythe site (Fig. 4), NIS richness remained consistently low throughout the sedimentary record, with only sporadic detections, such as *Neoporphyra haitanensis* and *Botrylloides violaceus* (Fig. 5). Similarly, at Bursledon, located at 5.5 km from the Hythe site, NIS richness remained similarly low and stable over time, with typically only a single non-indigenous species detected per year (Fig. 4). This consistent pattern across both sites is somewhat unexpected given the substantial shipping activity and industrial developments in Southampton Water since the mid-20^th^ century (Fig. S2), including the construction of container terminals and dockyards (Davis, 2012; Tavener, 1950). The observed peaks in Cu and Zn concentrations (Fig. S2), commonly linked to shipping infrastructure, motorway bridge construction during the 1970s (Bursledon), inputs from the Exxon oil refinery (Cu) and other manufacturing and production activity, and urban run-off, suggest periods of heightened industrial, port and urban activity (Appelt et al., (submitted); Boxall et al., 2000; Turner, 2010; Ytreberg et al., 2016). These trends are also consistent with historical shifts in antifouling paint formulations, which were predominantly Cu-based during the 1950s–1960s and increasingly incorporated Zn compounds as secondary biocides from the late 1960s onwards (Turner, 2010; Yebra et al., 2004; Ytreberg et al., 2016). However, these chemical signals of increased human activity did not coincide with a corresponding increase in NIS detections. This pattern suggests that, despite increased propagule pressure, additional ecological filters such as habitat unsuitability, competition, or pollutant stress may have constrained establishment success, underscoring the importance of local environmental context in mediating invasion outcomes.

One possible explanation for the absence of a recent increase in NIS richness is the potential limited resolution of both taxonomic identification and biogeographic assignment (Bik et al., 2012; Machida et al., 2017; Ratnasingham & Hebert, 2013; Zinger et al., 2019). A considerable proportion of ASVs could not be confidently assigned to species, across all centuries, limiting our ability to infer their likely origins throughout the record (Fig. S4). Even when taxonomy could be assigned, the lack of comprehensive invasion-history and population-genomic data for many estuarine taxa hindered confident classification as native, cryptogenic, or NIS (Rius & Turon, 2020). Consequently, several taxa were conservatively labelled as ‘unknown’. In contrast, native species richness exhibited a steady decline through time, particularly at Hythe, suggesting possible replacement dynamics (Fig. 4), habitat degradation, or competitive pressures from undetected or unclassified species introductions (Bishop et al., 2017; Halpern et al., 2015; López-Legentil et al., 2015).

The resulting temporal trajectories of GAMs revealed contrasting trends between the two cores (Fig. S5). At Hythe, species richness across all origin categories (native, cryptogenic, and non-indigenous species) declined gradually from the 19^th^ century onwards, consistent with the cumulative impacts of industrialisation, metal pollution, and urban development in the lower estuary (Appelt et al., (submitted); Halpern et al., 2015; Turner, 2010). In contrast, Bursledon, located further upstream, retained a stronger native signature until the late 20^th^ century, with only occasional detections of marine NIS across the core sequence. This spatial differentiation likely reflects the combined influence of salinity gradients, hydrodynamic conditions, and the distance from major shipping and industrial activity (Bishop et al., 2017; Johnston et al., 2017). These findings, although based on two sites, suggest that both anthropogenic disturbance and local environmental gradients jointly influence biodiversity trajectories through time (Cloern et al., 2016; DiBattista et al., 2022; Elliott & Whitfield, 2011; Kennish, 2002).

While these findings offer valuable insights into local-scale invasion histories, they likely represent only a subset of the estuarine mosaic. Broader spatial coverage, incorporating additional cores from inner and outer harbour zones, major port facilities, and less industrialised areas, could reveal more heterogeneous invasion patterns and a fuller picture of NIS dynamics across Southampton Water.

### Historical evidence of early species introductions

The detection of NIS as early as early as the 18^th^ century (Fig. 5), such as *N. vectensis*, a small estuarine sea anemone native to the east coast of North America, demonstrates that species introductions have been occurring for centuries (Darling et al., 2004; Reitzel et al., 2008). *N. vectensis* was likely introduced via early trans-Atlantic shipping routes, facilitated by Southampton’s growing importance as a trading port during the 1700s (Tavener, 1950).

Similarly, *N. haitanensis*, a red alga commonly cultivated in East Asia and transported for aquaculture, may have arrived also during the 18^th^ century through the movement of mariculture goods or hull fouling (Carlton, 1996). Although now dominant in some regions of the intertidal zone, the initial introduction of *N. haitanensis* went most likely unnoticed due to morphological similarity with native taxa and absence of molecular monitoring until recent decades (Chen et al., 2022). In contrast, the appearance of the solitary ascidian *S. plicata* (Fig. 5), a globally invasive tunicate with high tolerance to pollution and temperature fluctuations (Galià-Camps et al., 2023; Pineda et al., 2012; Platin & Shenkar, 2023), coincides with mid-20^th^-century industrial expansion and the opening of Southampton’s modern dock systems. The ecology of this ascidian suggests that contaminated, artificial substrates, such as ship hulls and harbour infrastructure, may have facilitated its settlement and spread in Southampton Water (Costello et al., 2022; Dafforn et al., 2011; Lambert, 2007; Paxton et al., 2025). In addition, localised pollution and metal enrichment, as indicated by XRF data and supported by previous geochemical studies (Appelt et al., (submitted)), may have excluded some sensitive benthic species in areas such as Hythe, where concentrations approach ecotoxicological thresholds, while levels across the wider estuary remained generally below toxicity limits (Sharifi et al., 1991). Such local environmental stressors likely created ecological space for tolerant taxa such as *S. plicata* to establish and persist where native competitors could not (Osborne & Poynton, 2019). As most marine benthic species have not evolved under polluted or industrially altered conditions, these changes may have provided novel ecological opportunities for pollution-tolerant invaders.

Another example of recent introductions detected in our sedaDNA record is *B. violaceus*, a colonial tunicate strongly associated with maritime traffic (Nelson, 2014; Wagstaff, 2017), was first detected in the uppermost sediment layers (Fig. 5). Its presence aligns with the opening of the container terminal 5 in 2014 in Southampton port and reflects increasing global shipping connectivity. This species is known for its capacity to rapidly colonise anthropogenic structures and for outcompeting native filter-feeders (Arenas et al., 2006; Lambert, 2007). Notably, *B. violaceus* was first reported in southern England during marina surveys in 2004 (Arenas et al., 2006), where it was already widespread along the south coast, including the Solent region. Subsequent surveys confirmed its persistence and patchy distribution across Solent marinas (Bishop et al., 2015), suggesting that the detection of this species in our sedaDNA data reflects a long-term presence of well-established populations.

The contrasting trends observed among NIS, some persisting for centuries, others appearing only transiently, and some expanding following recent industrialisation, highlight the importance of historical contingency and priority effects in community assembly and bioinvasion dynamics (Fukami, 2015; Kumschick & Richardson, 2013; Ricciardi et al., 2017; Stuble & Souza, 2016). The timing and sequence of species arrivals can affect whether non-indigenous taxa successfully establish, coexist with resident species, or fail to persist (Stuble & Souza, 2016; Torres et al., 2022). Although our sedaDNA record does not directly test these mechanisms, the contrasting temporal patterns observed hint that arrival history and environmental context likely interact to modulate invasion success over centennial timescales. Future multi-site sedaDNA comparisons could further explore how these processes jointly influence long-term community assembly and bioinvasion outcomes (Catford et al., 2009; Fukami, 2015).

### Limitations and methodological considerations

Our results on NIS dynamics emphasised the importance of taxonomic and biogeographic resolution for assessing invasion history. However, persistent blind spots in invasion science, particularly the incomplete representation of estuarine and benthic taxa in barcode reference databases, remain a major challenge for metabarcoding studies (Albano et al., 2021; Cowart et al., 2018; Katsanevakis et al., 2014; Weigand et al., 2019). Moreover, while the 18S marker provides broad eukaryotic coverage, its taxonomic resolution is generally lower than that of COI for many metazoans (Leray et al., 2013; Tang et al., 2012). Combining multiple genetic markers to maximise detection power and minimise misclassification is therefore essential (Stat et al., 2017; Taberlet et al., 2018). Addressing these limitations through improved reference databases and the establishment of long-term baseline invasion records will be critical to more precisely resolve species’ origins and to understand the role of NIS in shaping biodiversity change (Ficetola et al., 2018).

While our analytical framework included extensive technical replication and stringent filtering to minimise false positives, our analyses may have excluded biologically valid but low-abundant taxa (Lanzén et al., 2017). Likewise, metabarcoding read abundance is influenced by stochastic amplification, primer bias, and DNA degradation, and thus should not be interpreted as a direct proxy for organismal biomass (Deiner et al., 2017; Elbrecht & Leese, 2015). Addressing these limitations will require experimental calibration, mock communities, and quantitative assays to refine detection thresholds and improve the ecological interpretability of sedaDNA signals (Zinger et al., 2019).

Future studies should expand the use of shotgun metagenomics and multi-marker approaches to enhance taxonomic and functional resolution, particularly for underrepresented groups such as soft-bodied invertebrates (Stat et al., 2017; Taberlet et al., 2018). Parallel efforts to curate regional reference libraries and integrate genetic data with ecological and functional traits will also improve our biogeographic status assignments and more accurately assess invasion processes (Creer et al., 2016; Weigand et al., 2019).

### Historical insights and implications for biosecurity

Our findings support the need to extend biosecurity baselines further into the past. The detection of historical introductions predating modern port infrastructure, already occurring in the 18^th^ century, challenges prevailing assumptions about the onset of marine invasions and highlights the importance of incorporating pre-industrial conditions into restoration and conservation targets (Campbell et al., 2025; Holman et al., 2025a; Ojaveer et al., 2018). Furthermore, integrating sedaDNA with complementary palaeoenvironmental proxies such as lipid biomarkers, isotopic analyses, and sediment chemistry offers a powerful means of reconstructing the mechanisms underlying past biological invasions (Capo et al., 2021; Coolen et al., 2013).

We initially hypothesised that intensified maritime activity during the mid-20^th^ century contributed to a rise in NIS richness and significantly altered community composition. In addition, we expected trace metal (Cu, Zn, and Pb) concentrations to reflect industrial and shipping pressures that might correlate with community shifts. However, our results revealed more complex temporal patterns. Despite clear industrial signals in the sediment record, NIS richness remained stable, suggesting that invasion outcomes are shaped by a multifactorial interplay of propagule pressure, environmental filters, and historical contingencies.

### Conclusions

By bridging molecular palaeoecology and invasion science, our study provides new insights into the temporal dynamics of NIS, including potential failed introductions, and highlights the intrinsic characteristics of the ongoing transformation of estuarine ecosystems under sustained human influence.

Our work demonstrates the value of intertidal sedaDNA archives for reconstructing centuries of ecological change, offering a framework with broad applicability for biosecurity, conservation baselines and long-term biodiversity assessments.

## Author contributions

Andy Cundy, Luke Holman, Nicola Pratt and Marc Rius conducted the field sampling. Nicola Pratt conducted the core subsampling and DNA extraction. Luke Holman undertook the molecular laboratory work. Sandra Nogué and Andy Cundy were responsible for sediment dating, and geochemical and palaeoecological analyses. Elena Baños carried out laboratory and bioinformatic analyses, data interpretation, statistical analyses, and led manuscript preparation. All authors contributed to manuscript revision and approved the final version for submission.

## Data Availability Statement

Raw sequence reads are deposited in the NCBI Sequence Read Archive under BioProject PRJNA1364850. All additional processed and supplementary data supporting this study have been permanently archived under DOI: 10.5281/zenodo.17647494.

## Funding Statement

This work was supported by a SMMI Research Collaboration Stimulus Fund grant entitled “A palaeoecological approach to monitor invasive species” from the University of Southampton, the Spanish Ministry of Science, Innovation and Universities grant PID2020-118550RB (MICIU/AEI/https://doi.org/10.13039/501100011033), the grant TED2021-132228B-C22 funded by MCIN/AEI/10.13039/501100011033 and the European Union (“NextGenerationEU”/PRTR); and the Spanish Ministry of Science, Innovation and Universities grant PID2023-146307OB-C22 (MICIU/AEI/10.13039/501100011033 and FEDER, UE).

## Conflict of Interest Disclosure

The authors declare no conflict of interest.

## Ethics Approval Statement

Not applicable. This study did not involve experiments on live animals, humans, or endangered species.

## Patient Consent Statement

Not applicable.

## Permission to Reproduce Material from Other Sources

Not applicable. All figures and data presented are original to this study.

## Clinical Trial Registration

Not applicable.

## Acknowledgements

This work was supported by a SMMI Research Collaboration Stimulus Fund grant entitled “A palaeoecological approach to monitor invasive species” from the University of Southampton, the Spanish Ministry of Science, Innovation and Universities grant PID2020-118550RB (MICIU/AEI/https://doi.org/10.13039/501100011033), the grant TED2021-132228B-C22 funded by MCIN/AEI/10.13039/501100011033 and the European Union (“NextGenerationEU”/PRTR); the British Ocean Sediment Core Research Facility (BOSCORF, National Oceanography Centre, Southampton); and the Spanish Ministry of Science, Innovation and Universities grant PID2023-146307OB-C22 (MICIU/AEI/10.13039/501100011033 and FEDER, UE). Dr. David Reading (GAU-Radioanalytical, University of Southampton) is thanked for ^210^Pb and ^137^Cs analytical support.

## Data Accessibility and Benefit-Sharing

### Data Accessibility Statement

Raw sequence reads are deposited in the NCBI Sequence Read Archive under BioProject PRJXXXXXX. All additional processed and supplementary data supporting this study have been permanently archived under DOI: 10.5281/zenodo.17474797.

### Benefit-Sharing Statement

Benefits from this research accrue from the open sharing of all generated data and results through public databases, as described above. The work was carried out in collaboration between UK- and Spain-based institutions, with all contributing scientists included as co-authors. The study addresses priority concerns in marine biodiversity conservation, including the reconstruction of invasion histories and the identification of environmental drivers influencing non-indigenous species establishment.

## Supplementary Information

**Supplementary Data S1:** X-Ray Fluorescens and Dating information.

**Supplementary Data S2.1:** Total abundance matrix of filtered amplicon sequence variants (ASVs) obtained from COI metabarcoding of sedimentary ancient DNA (sedaDNA) samples from the Hythe (HM01, HM03) and Bursledon (HB5) cores. Values represent the total number of reads assigned to each ASV per sample after quality filtering and curation.

**Supplementary Data S2.2:** Total abundance matrix of filtered amplicon sequence variants (ASVs) obtained from 18S metabarcoding of sedimentary ancient DNA (sedaDNA) samples from the Hythe (HM01, HM03) and Bursledon (HB5) cores. Values represent the total number of reads assigned to each ASV per sample after quality filtering and curation.

**Supplementary Data S3:** Relative abundance matrices of filtered amplicon sequence variants (ASVs) for both COI and 18S metabarcoding datasets. Values represent the proportional read abundance of each ASV per sample after normalisation, showing community composition changes across sediment cores and depths.

**Supplementary Data S4.** List of detected species classified by their status: NAT (native species), CRY (cryptogenic species), NIS (non-indigenous species) and UNK (taxa with unknown status).

**Table S1.** Summary of core data, including ID, total length, and description of each core.

| Core ID | UTM coordinates | Depth (cm) | Description |
| --- | --- | --- | --- |
| HM01 | 619883.1, 5638889.9 | 50,8 | The core consists entirely of muddy sediment. The top 20 cm contains alternating thin layers of brown-orange and grey muds. Below this, a 10 cm layer of dark brown mud is present, followed by the bottom 20 cm, which is composed of dark grey mud. |
| HM03 | 613427.4, 5635923.5 | 149,8 | The core consists entirely of grey muddy sediment. |
| HB5 | 613499.9, 5635915.9 | 50 | The core consists entirely of muddy sediment. The top 15 cm contains alternating thin layers of brown-orange and grey fine sediment. A 7 cm layer of dark brown mud occurs below this. The basal 35 cm of the core is composed of dark grey mud. |

**Table S2a.** **a.** General Linear Models comparing genus richness between Hythe and Bursledon sites for COI and 18S genes.

|  |  | Sites | Estimate | SE | df | T | p-value |
| --- | --- | --- | --- | --- | --- | --- | --- |
| Richness | COI | Hythe -<br>Bursledon | 1.59 | 0.38 | 526 | 4.174 | <.0001 |
|  | 18S | Hythe -<br>Bursledon | 13.5 | 1.83 | 524 | 7.371 | <.0001 |

**Table S2b.**
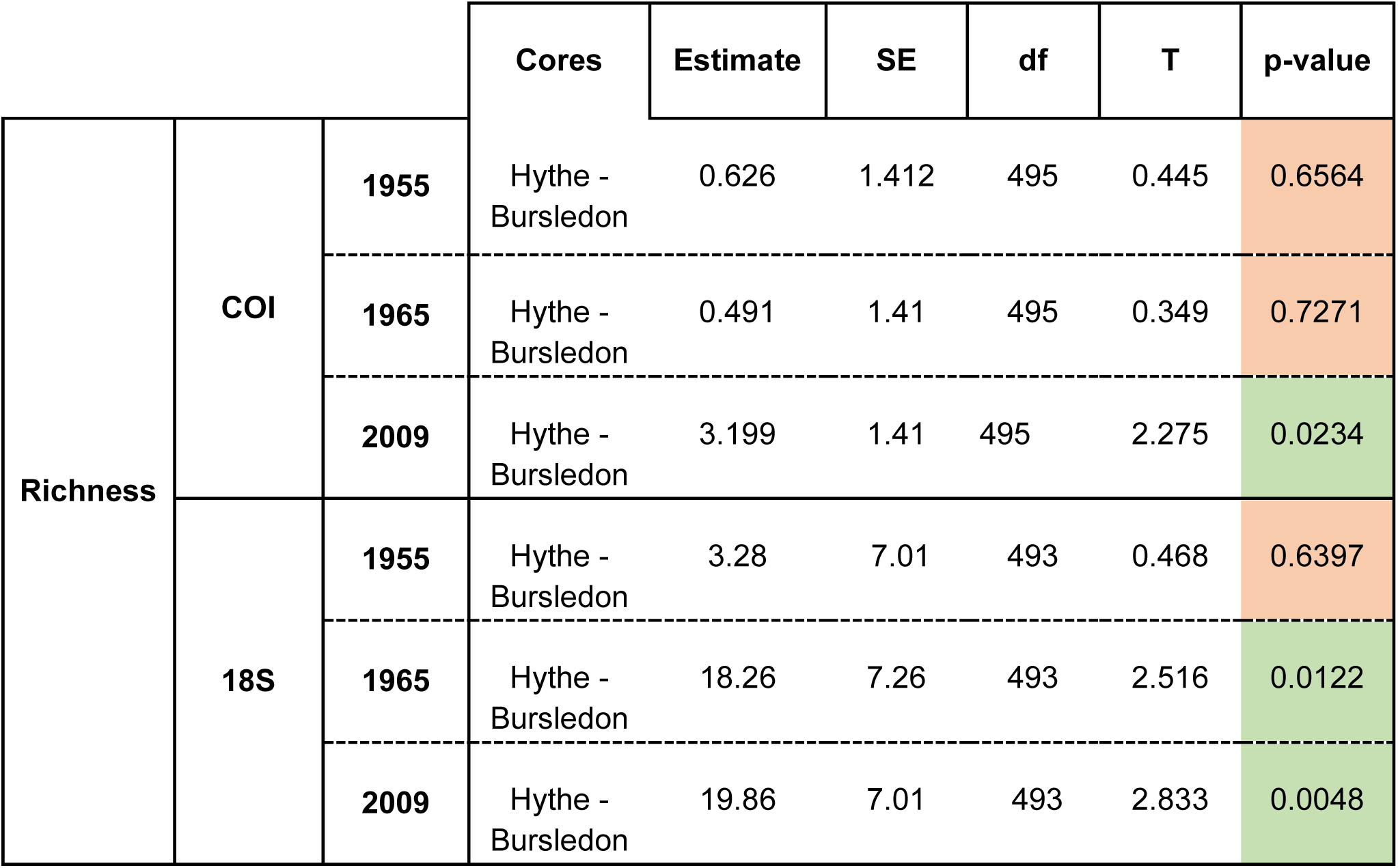
General Linear Models testing differences in genus richness between Hythe and Bursledon across three representative years (1955, 1965, 2009).

**Table S3.** Generalised Additive Model results with effective degrees of freedom (edf), F-tests (F) and p-values.

|  |  | Cores | edf | F | p-value |
| --- | --- | --- | --- | --- | --- |
| GAM | Hythe | NIS | 1 | 2.868 | 0.0940 |
|  |  | NAT | 1 | 57.735 | <2e-16 |
|  |  | CRY | 1 | 0.001 | 0.9721 |
|  |  | Unknown | 1 | 25.529 | <6.6e-06 |
| GAM | Bursledon | NIS | 1 | 0.058 | 0.8181 |
|  |  | NAT | 1 | 0.079 | 0.7781 |
|  |  | CRY | 1 | 0.149 | 0.7004 |
|  |  | Unknown | 1 | 5.963 | 0.0167 |

**Table S4:**
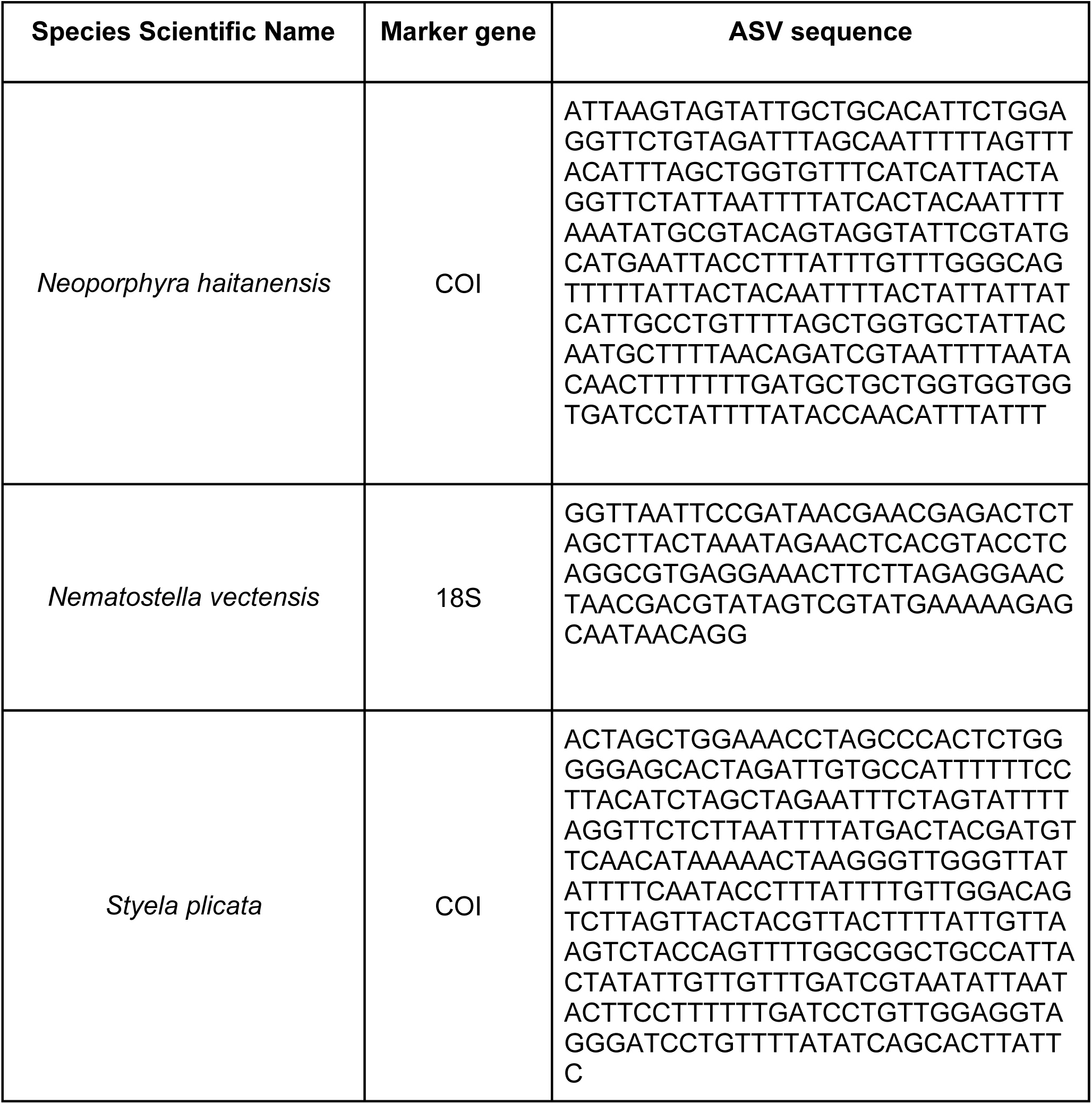
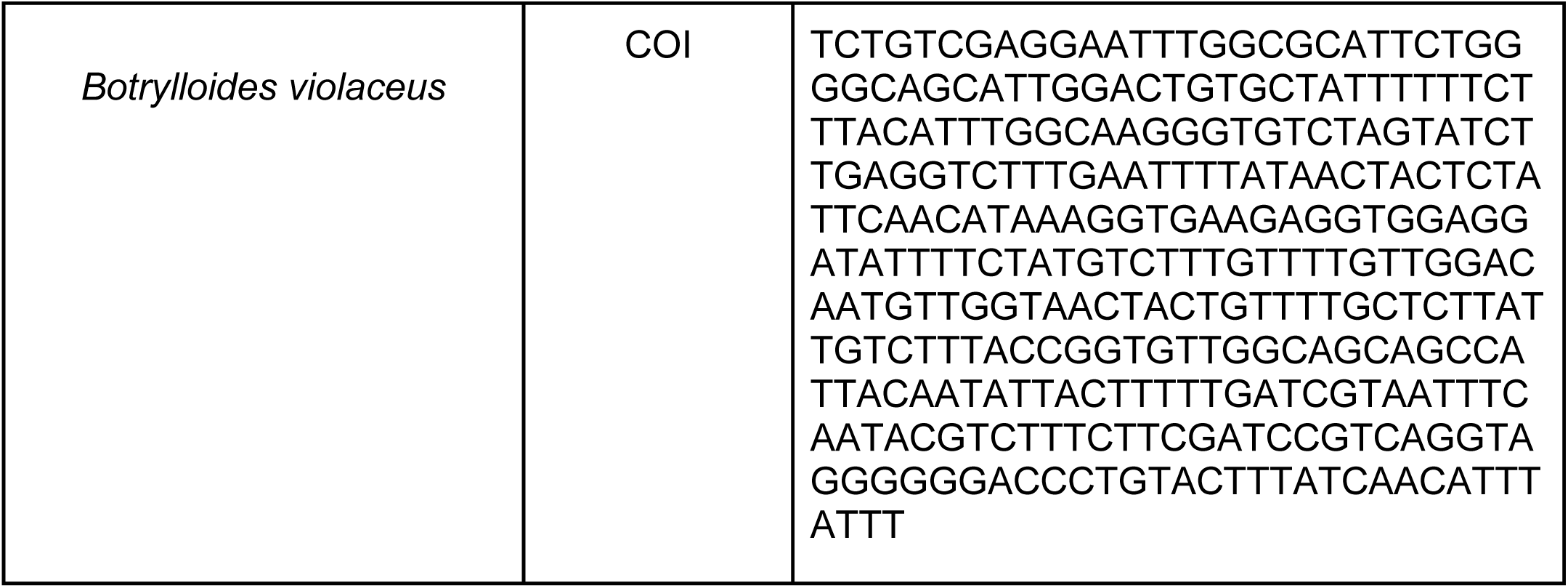
Raw amplicon sequence variants (ASVs) corresponding to the key non-indigenous and native species detected in this study. For each taxon, the marker gene (COI or 18S) and the representative ASV sequence retained after quality filtering and LULU curation are shown. These sequences represent the exact barcode variants recovered from sedimentary ancient DNA (sedaDNA) analyses and were used to confirm species-level identifications of *Neoporphyra haitanensis*, *Nematostella vectensis*, *Styela plicata*, and *Botrylloides violaceus*.

**Fig. S1.**
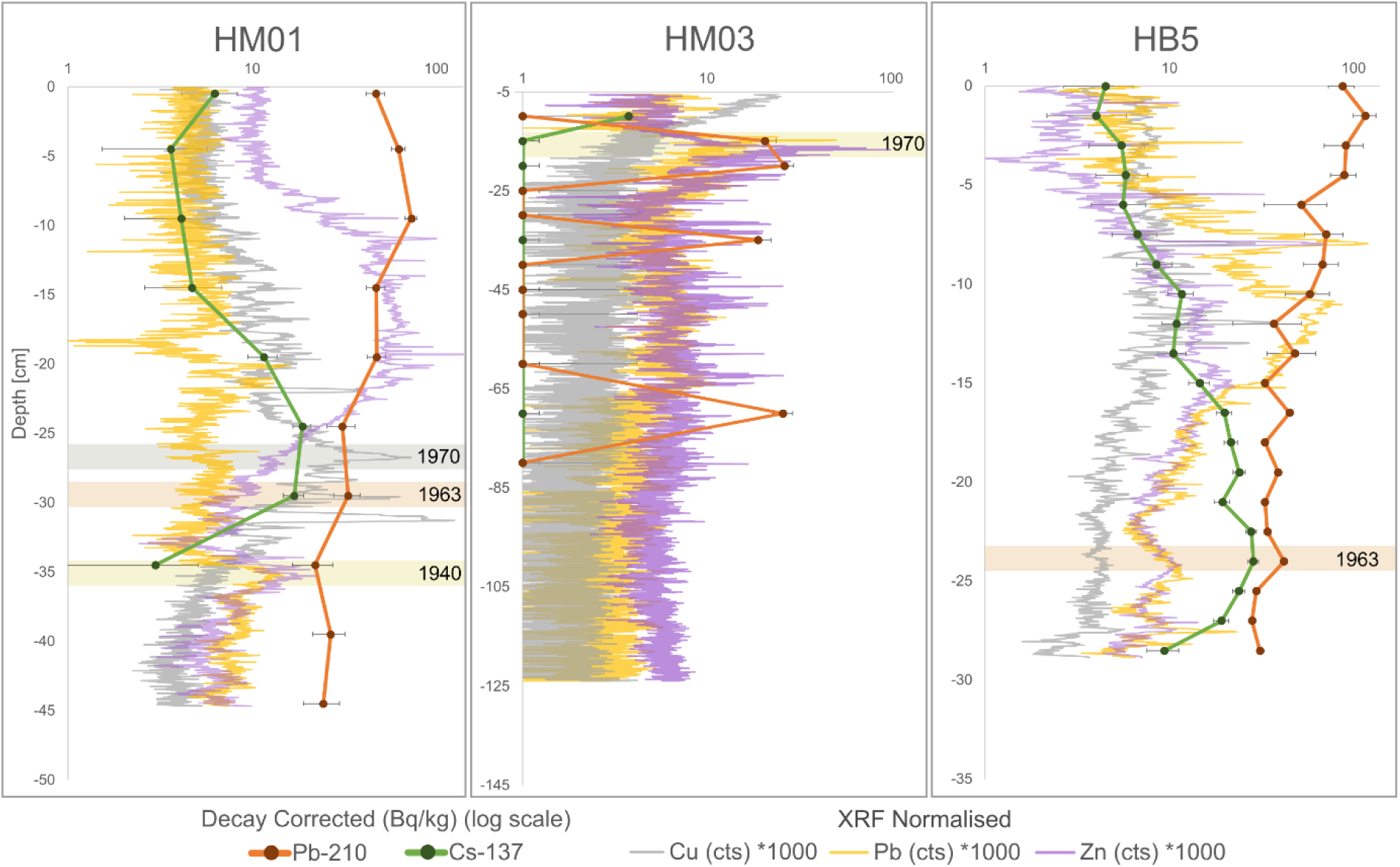
^210^Pb and ^137^Cs activities, and scanning XRF data for Cu, Pb and Zn (counts (cts) normalised to total scatter) with core depth. . Dates are based on ^210^Pb and ^137^Cs-derived sediment accumulation rates, and known metal input maxima (see text for discussion).

**Fig. S2:**
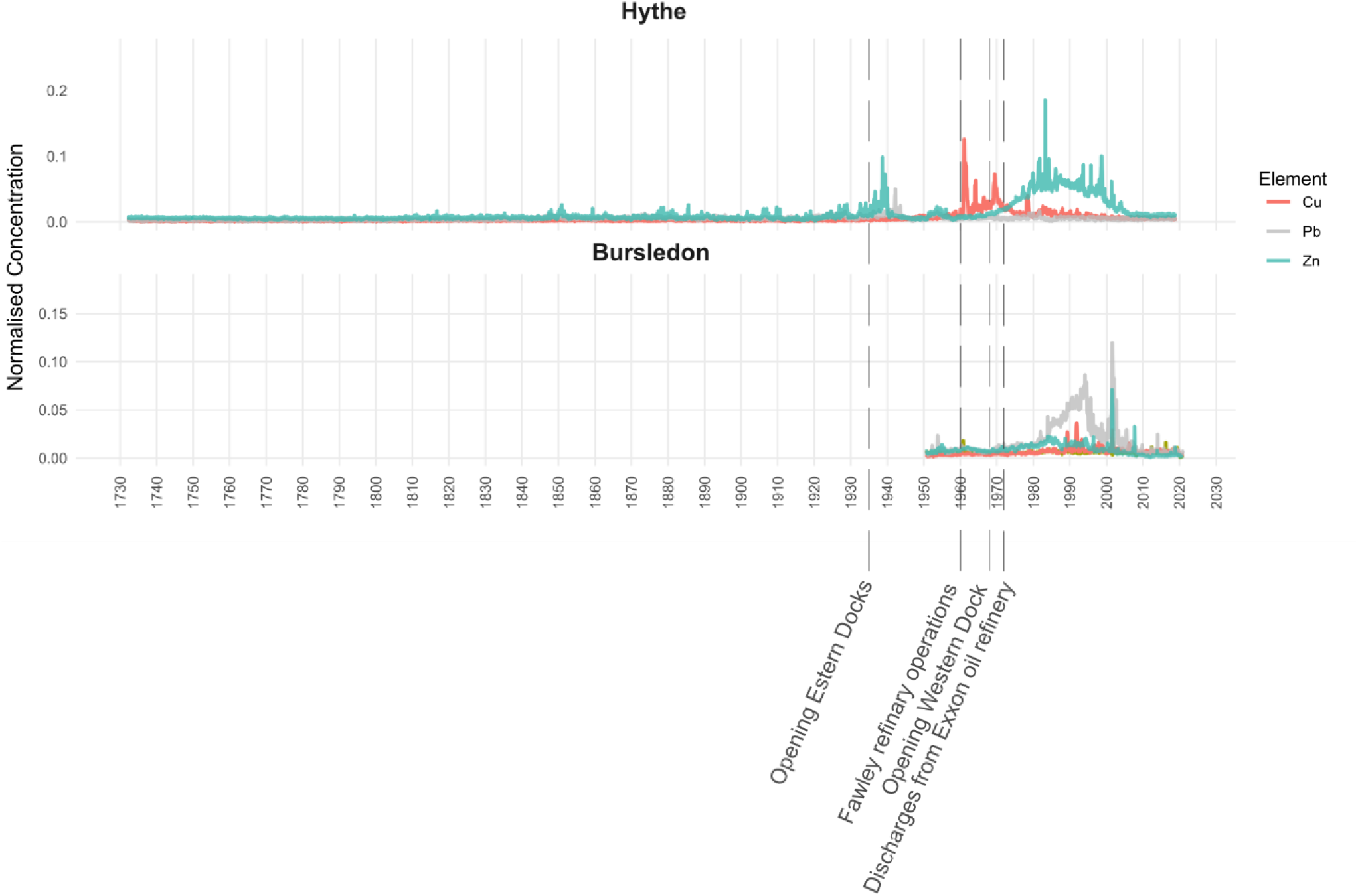
Normalised concentration of Zn, Cu, Pb and Ca measured by scanning XRF in the two dated sediment cores from Southampton Water. The profiles show a marked increase in Zn, Cu and Pb concentrations from the mid-19^th^ century onwards, reflecting the onset of industrial activity and the influence of antifouling paint residues.

**Fig. S3:**
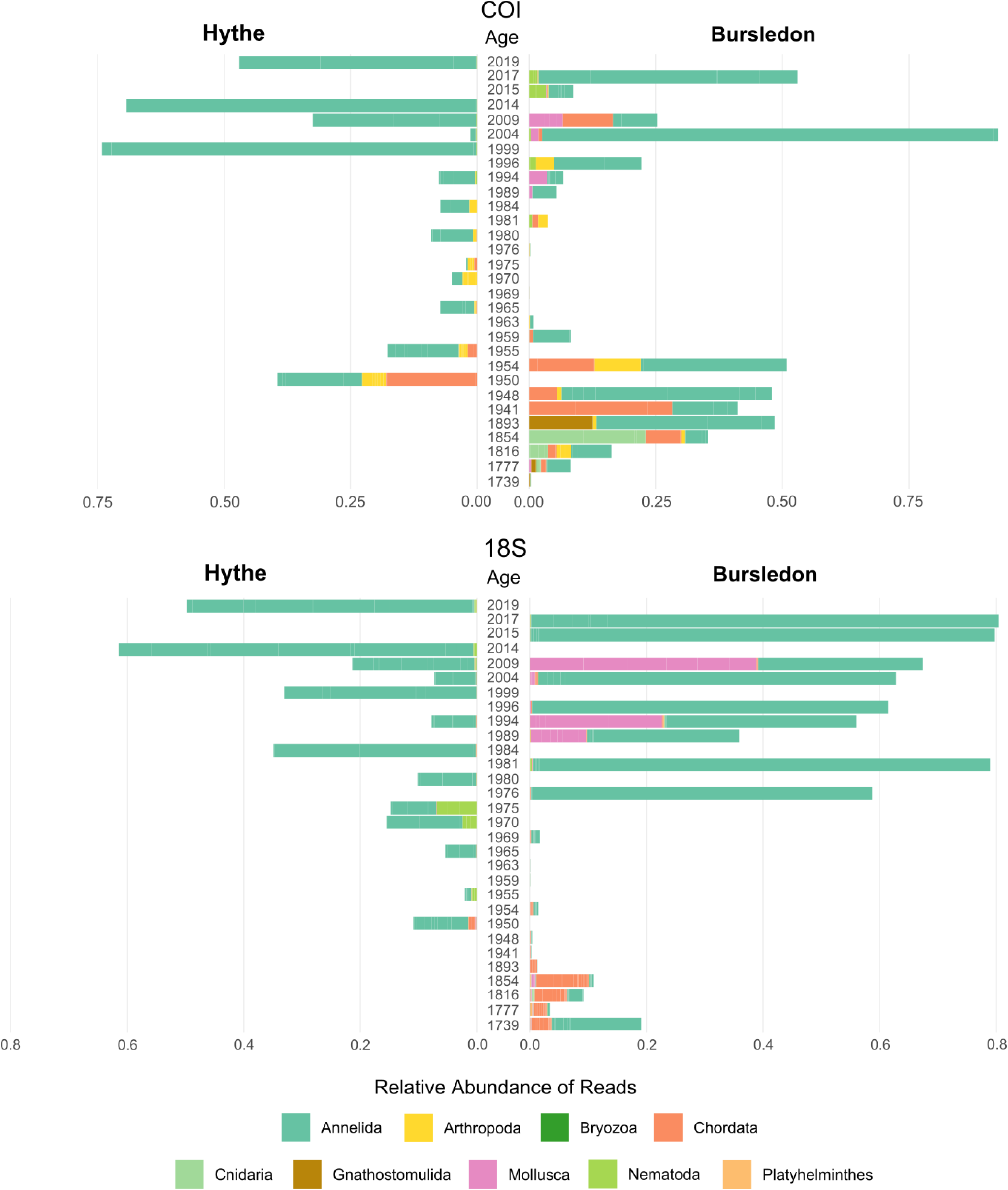
BarPlot of relative abundance of metazoan phyla across dated sediment layers for the COI and 18S datasets respectively.

**Fig. S4:**
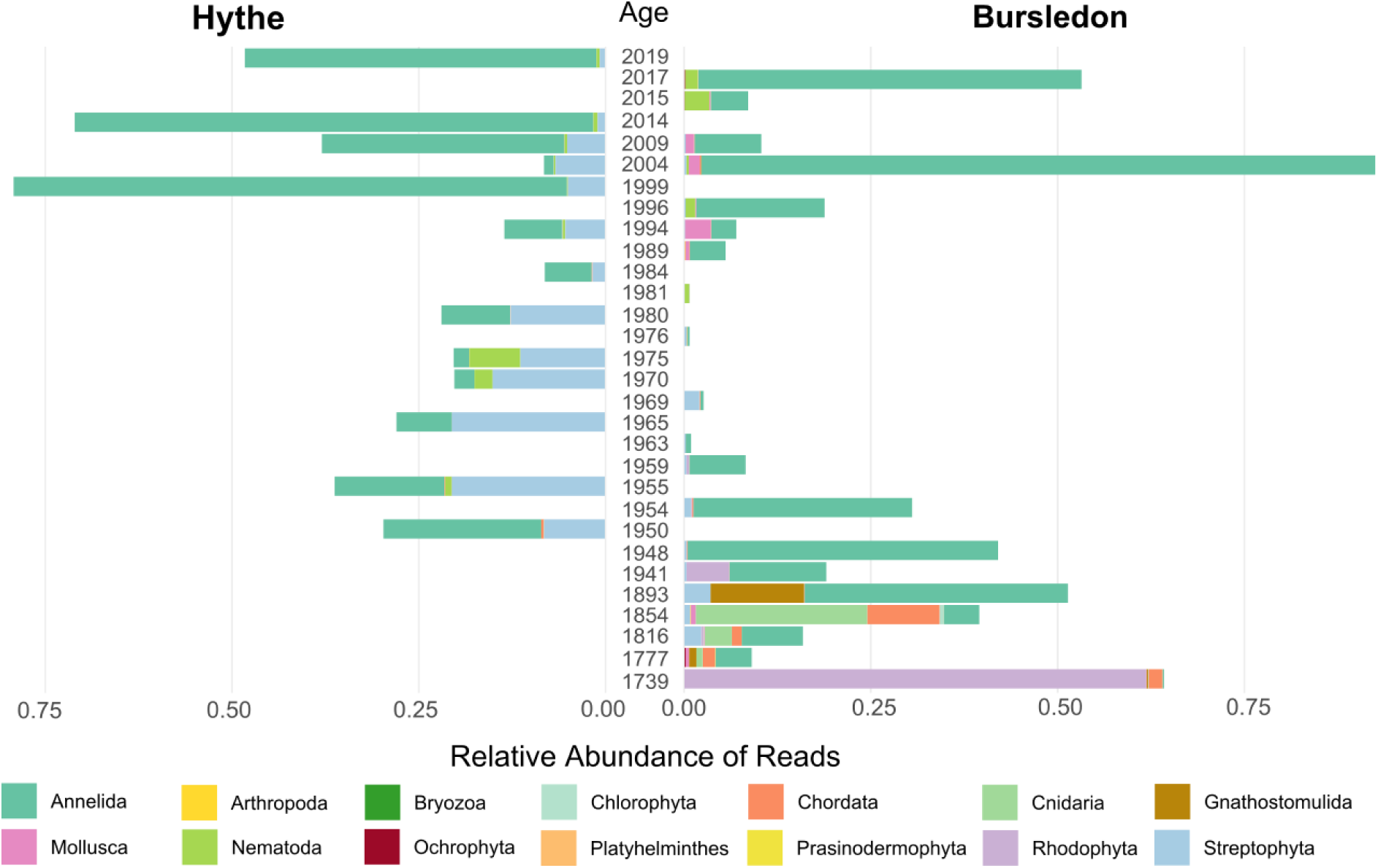
Composition of identified and status classified Phyla over time in the studied sites.

**Fig. S5:**
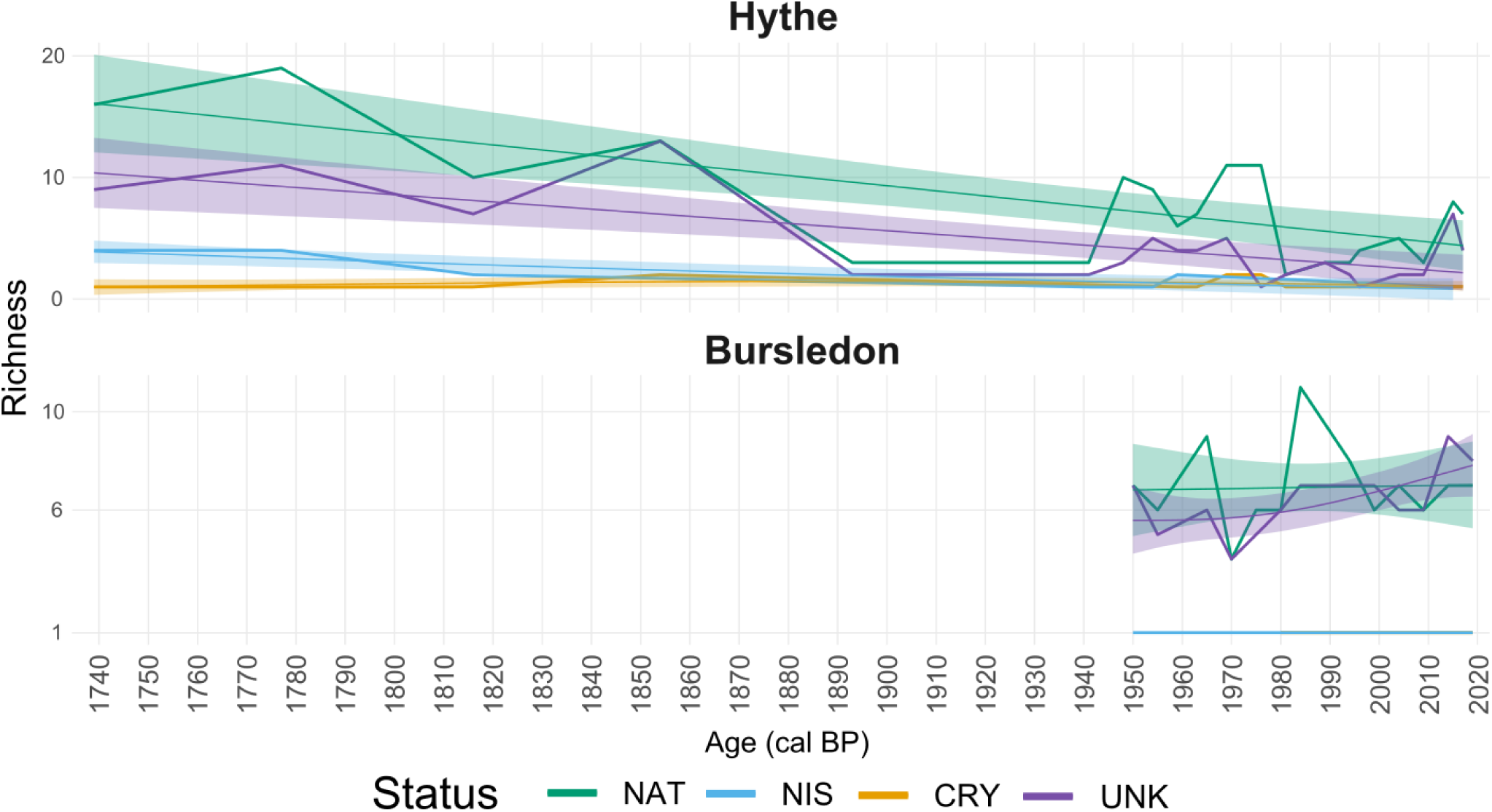
Temporal trends in species richness by biogeographic status in Hythe and Bursledon cores. Generalised Additive Model (GAM) curves represent the richness of classified species over time for Non-Indigenous Species (NIS, blue), Native (NAT, green), and Cryptogenic (CRY, red) taxa. Shaded areas indicate 95% confidence intervals. Note that the x-axis in the Bursledon panel starts at 1950 due to the sediment record range in that core.

## Notes

https://doi.org/10.5281/zenodo.17647494

### Competing Interest Statement

The authors have declared no competing interest.

## References

Albano, P. G., Steger, J., Bošnjak, M., Dunne, B., Guifarro, Z., Turapova, E., Hua, Q., Kaufman, D. S., Rilov, G., & Zuschin, M. (2021). Native biodiversity collapse in the eastern Mediterranean. Proceedings of the Royal Society B: Biological Sciences, 288(1942), 20202469. 10.1098/rspb.2020.2469

Alsos, I. G., Boussange, V., Rijal, D. P., Beaulieu, M., Brown, A. G., Herzschuh, U., Svenning, J.-C., & Pallissier, L. (2023, November 7). Ancient sedimentary DNA to forecast trajectories of ecosystem under climate change [NonPeerReviewed]. https://eprints.soton.ac.uk/488059/

Appleby, P. G. (2002). Chronostratigraphic Techniques in Recent Sediments. In W. M. Last & J. P. Smol (Eds.), Tracking Environmental Change Using Lake Sediments (Vol. 1, pp. 171–203). Kluwer Academic Publishers. 10.1007/0-306-47669-X_9

Appleby, P. G., & Oldfield, F. (1978). The calculation of lead-210 dates assuming a constant rate of supply of unsupported ^210^Pb to the sediment. CATENA, 5(1), 1–8. 10.1016/S0341-8162(78)80002-2

Appelt, J.-S., Cundy, A., Whiteside, J., & Bray, P. (submitted). Spatial and Temporal Trends in Trace Element, PAH and Steroidal Hormone Contamination in a Major Urban and Industrial Estuary: Southampton Water, UK. Environmental Science and Technology.

Arenas, F., Bishop, J. D. D., Carlton, J. T., Dyrynda, P. J., Farnham, W. F., Gonzalez, D. J., Jacobs, M. W., Lambert, C., Lambert, G., Nielsen, S. E., Pederson, J. A., Porter, J. S., Ward, S., & Wood, C. A. (2006). Alien species and other notable records from a rapid assessment survey of marinas on the south coast of England. Journal of the Marine Biological Association of the United Kingdom, 86(6), 1329–1337. 10.1017/S0025315406014354

Armbrecht, L. H. (2020). The Potential of Sedimentary Ancient DNA to Reconstruct Past Ocean Ecosystems. Oceanography, 33(2), 116–123.

Armbrecht, L., Hallegraeff, G., Bolch, C. J. S., Woodward, C., & Cooper, A. (2021). Hybridisation capture allows DNA damage analysis of ancient marine eukaryotes. Scientific Reports, 11(1), 3220. 10.1038/s41598-021-82578-6

Bailey, S. A., Brown, L., Campbell, M. L., Canning-Clode, J., Carlton, J. T., Castro, N., Chainho, P., Chan, F. T., Creed, J. C., Curd, A., Darling, J., Fofonoff, P., Galil, B. S., Hewitt, C. L., Inglis, G. J., Keith, I., Mandrak, N. E., Marchini, A., McKenzie, C. H., … Zhan, A. (2020). Trends in the detection of aquatic non-indigenous species across global marine, estuarine and freshwater ecosystems: A 50-year perspective. Diversity and Distributions, 26(12), 1780–1797. 10.1111/ddi.13167

Bik, H. M., Porazinska, D. L., Creer, S., Caporaso, J. G., Knight, R., & Thomas, W. K. (2012). Sequencing our way towards understanding global eukaryotic biodiversity. Trends in Ecology & Evolution, 27(4), 233–243. 10.1016/j.tree.2011.11.010

Bishop, J., Wood, C., Yunnie, A., & Griffiths, C. (2015). Unheralded arrivals: Non-native sessile invertebrates in marinas on the English coast. Aquatic Invasions, 10(3), 249–264. 10.3391/ai.2015.10.3.01

Bishop, M. J., Mayer-Pinto, M., Airoldi, L., Firth, L. B., Morris, R. L., Loke, L. H. L., Hawkins, S. J., Naylor, L. A., Coleman, R. A., Chee, S. Y., & Dafforn, K. A. (2017). Effects of ocean sprawl on ecological connectivity: Impacts and solutions. Journal of Experimental Marine Biology and Ecology, 492, 7–30. 10.1016/j.jembe.2017.01.021

Bodur, S. O., Samuel, S. O., Polat, M. F., Aycan, M., & Asiloglu, R. (2025). Protists exhibit community-level adaptation and functional redundancy under gradient soil salinity. Science of The Total Environment, 981, 179606. 10.1016/j.scitotenv.2025.179606

Boxall, A. B. A., Comber, S. D., Conrad, A. U., Howcroft, J., & Zaman, N. (2000). Inputs, Monitoring and Fate Modelling of Antifouling Biocides in UK Estuaries. Marine Pollution Bulletin, 40(11), 898–905. 10.1016/S0025-326X(00)00021-7

Callahan, B. J., McMurdie, P. J., Rosen, M. J., Han, A. W., Johnson, A. J. A., & Holmes, S. P. (2016). DADA2: High-resolution sample inference from Illumina amplicon data. Nature Methods, 13(7), 581–583. 10.1038/nmeth.3869

Camacho, C., Coulouris, G., Avagyan, V., Ma, N., Papadopoulos, J., Bealer, K., & Madden, T. L. (2009). BLAST+: Architecture and applications. BMC Bioinformatics, 10, 421. 10.1186/1471-2105-10-421

Campbell, M. A., Ward, I., Blyth, A., & Allentoft, M. E. (2025). Using sedimentary ancient DNA in coastal and marine contexts to explore past human–environmental interactions in Australia. Philosophical Transactions of the Royal Society B: Biological Sciences, 380(1930), 20240032. 10.1098/rstb.2024.0032

Capo, E., Giguet-Covex, C., Rouillard, A., Nota, K., Heintzman, P. D., Vuillemin, A., Ariztegui, D., Arnaud, F., Belle, S., Bertilsson, S., Bigler, C., Bindler, R., Brown, A. G., Clarke, C. L., Crump, S. E., Debroas, D., Englund, G., Ficetola, G. F., Garner, R. E., … Parducci, L. (2021). Lake Sedimentary DNA Research on Past Terrestrial and Aquatic Biodiversity: Overview and Recommendations. Quaternary, 4(1), Article 1. 10.3390/quat4010006

Carlton, J. T. (1996). Marine Bioinvasions: The Alteration of Marine Ecosystems by Nonindigenous Species. Oceanography, 9(1), 36–43.

Carlton, J. T., & Schwindt, E. (2024). The assessment of marine bioinvasion diversity and history. Biological Invasions, 26(1), 237–298. 10.1007/s10530-023-03172-7

Catford, J. A., Jansson, R., & Nilsson, C. (2009). Reducing redundancy in invasion ecology by integrating hypotheses into a single theoretical framework. Diversity and Distributions, 15(1), 22–40. 10.1111/j.1472-4642.2008.00521.x

Celis-Hernandez, O., Cundy, A. B., Croudace, I. W., & Ward, R. D. (2022). Environmental risk of trace metals and metalloids in estuarine sediments: An example from Southampton Water, U.K. Marine Pollution Bulletin, 178, 113580. 10.1016/j.marpolbul.2022.113580

Chen, H., Chu, J. S.-C., Chen, J., Luo, Q., Wang, H., Lu, R., Zhu, Z., Yuan, G., Yi, X., Mao, Y., Lu, C., Wang, Z., Gu, D., Jin, Z., Zhang, C., Weng, Z., Li, S., Yan, X., & Yang, R. (2022). Insights into the Ancient Adaptation to Intertidal Environments by Red Algae Based on a Genomic and Multiomics Investigation of Neoporphyra haitanensis. Molecular Biology and Evolution, 39(1), msab315. 10.1093/molbev/msab315

Cloern, J. E., Abreu, P. C., Carstensen, J., Chauvaud, L., Elmgren, R., Grall, J., Greening, H., Johansson, J. O. R., Kahru, M., Sherwood, E. T., Xu, J., & Yin, K. (2016). Human activities and climate variability drive fast-paced change across the world’s estuarine–coastal ecosystems. Global Change Biology, 22(2), 513–529. 10.1111/gcb.13059

Collins, R. A., Wangensteen, O. S., O’Gorman, E. J., Mariani, S., Sims, D. W., & Genner, M. J. (2018). Persistence of environmental DNA in marine systems. Communications Biology, 1(1), 185. 10.1038/s42003-018-0192-6

Coolen, M. J. L., Orsi, W. D., Balkema, C., Quince, C., Harris, K., Sylva, S. P., Filipova-Marinova, M., & Giosan, L. (2013). Evolution of the plankton paleome in the Black Sea from the Deglacial to Anthropocene. Proceedings of the National Academy of Sciences, 110(21), 8609–8614. 10.1073/pnas.1219283110

Costello, K. E., Lynch, S. A., McAllen, R., O’Riordan, R. M., & Culloty, S. C. (2022). Assessing the potential for invasive species introductions and secondary spread using vessel movements in maritime ports. Marine Pollution Bulletin, 177, 113496. 10.1016/j.marpolbul.2022.113496

Cowart, D. A., Murphy, K. R., & Cheng, C.-H. C. (2018). Metagenomic sequencing of environmental DNA reveals marine faunal assemblages from the West Antarctic Peninsula. Marine Genomics, 37, 148–160. 10.1016/j.margen.2017.11.003

Creer, S., Deiner, K., Frey, S., Porazinska, D., Taberlet, P., Thomas, W. K., Potter, C., & Bik, H. M. (2016). The ecologist’s field guide to sequence-based identification of biodiversity. Methods in Ecology and Evolution, 7(9), 1008–1018. 10.1111/2041-210X.12574

Croudace, I. W., & Cundy, A. B. (1995). Heavy Metal and Hydrocarbon Pollution in Recent Sediments from Southampton Water, Southern England: A Geochemical and Isotopic Study. Environmental Science & Technology, 29(5), 1288–1296. 10.1021/es00005a021

Croudace, I. W., Löwemark, L., Tjallingii, R., & Zolitschka, B. (2019). High resolution XRF core scanners: A key tool for the environmental and palaeoclimate sciences. Quaternary International, 514, 1–4. 10.1016/j.quaint.2019.05.038

Cundy, A. B., & Croudace, I. W. (2017). The Fate of Contaminants and Stable Pb Isotopes in a Changing Estuarine Environment: 20 Years On. Environmental Science & Technology, 51(17), 9488–9497. 10.1021/acs.est.7b00973

Cundy, A. B., Croudace, I. W., Cearreta, A., & Irabien, M. J. (2003). Reconstructing historical trends in metal input in heavily-disturbed, contaminated estuaries: Studies from Bilbao, Southampton Water and Sicily. Applied Geochemistry, 18(2), 311–325. 10.1016/S0883-2927(02)00127-0

Cundy, A. B., Croudace, I. W., Thomson, J., & Lewis, J. T. (1997). Reliability of Salt Marshes as “Geochemical Recorders” of Pollution Input: A Case Study from Contrasting Estuaries in Southern England. Environmental Science & Technology, 31(4), 1093–1101. 10.1021/es960622d

Dafforn, K. A., Lewis, J. A., & Johnston, E. L. (2011). Antifouling strategies: History and regulation, ecological impacts and mitigation. Marine Pollution Bulletin, 62(3), 453–465. 10.1016/j.marpolbul.2011.01.012

Darling, J. A., & Carlton, J. T. (2018). A Framework for Understanding Marine Cosmopolitanism in the Anthropocene. Frontiers in Marine Science, 5. 10.3389/fmars.2018.00293

Darling, J., Reitzel, A., & Finnerty, J. (2004). Regional population structure of a widely introduced estuarine invertebrate: Nematostella vectensis Stephenson in New England. Molecular Ecology, 13, 2969–2981. 10.1111/j.1365-294X.2004.02313.x

Davis, R. (2012). The Rise of the English Shipping Industry in the Seventeenth and Eighteenth Centuries. Oxford University Press.

Deiner, K., Bik, H. M., Mächler, E., Seymour, M., Lacoursière-Roussel, A., Altermatt, F., Creer, S., Bista, I., Lodge, D. M., de Vere, N., Pfrender, M. E., & Bernatchez, L. (2017). Environmental DNA metabarcoding: Transforming how we survey animal and plant communities. Molecular Ecology, 26(21), 5872–5895. 10.1111/mec.14350

DiBattista, J. D., Fowler, A. M., Riley, I. J., Reader, S., Hay, A., Parkinson, K., & Hobbs, J.-P. A. (2022). The use of environmental DNA to monitor impacted coastal estuaries. Marine Pollution Bulletin, 181, 113860. 10.1016/j.marpolbul.2022.113860

Domaizon, I., Winegardner, A., Capo, E., Gauthier, J., & Gregory-Eaves, I. (2017). DNA-based methods in paleolimnology: New opportunities for investigating long-term dynamics of lacustrine biodiversity. Journal of Paleolimnology, 58(1), 1–21. 10.1007/s10933-017-9958-y

Dornelas, M., Gotelli, N. J., McGill, B., Shimadzu, H., Moyes, F., Sievers, C., & Magurran, A. E. (2014). Assemblage Time Series Reveal Biodiversity Change but Not Systematic Loss. Science, 344(6181), 296–299. 10.1126/science.1248484

Elbrecht, V., & Leese, F. (2015). Can DNA-Based Ecosystem Assessments Quantify Species Abundance? Testing Primer Bias and Biomass—Sequence Relationships with an Innovative Metabarcoding Protocol. PLOS ONE, 10(7), e0130324. 10.1371/journal.pone.0130324

Elliott, M., & Whitfield, A. K. (2011). Challenging paradigms in estuarine ecology and management. Estuarine, Coastal and Shelf Science, 94(4), 306–314. 10.1016/j.ecss.2011.06.016

Ficetola, G. F., Miaud, C., Pompanon, F., & Taberlet, P. (2008). Species detection using environmental DNA from water samples. Biol. Lett., 4423–425. 10.1098/rsbl.2008.0118

Ficetola, G. F., Poulenard, J., Sabatier, P., Messager, E., Gielly, L., Leloup, A., Etienne, D., Bakke, J., Malet, E., Fanget, B., Støren, E., Reyss, J.-L., Taberlet, P., & Arnaud, F. (2018). DNA from lake sediments reveals long-term ecosystem changes after a biological invasion. Science Advances, 4(5), eaar4292. 10.1126/sciadv.aar4292

Fisher, S., Goff, J., Cundy, A. B., & Sear, D. (2025). The tsunami history and prehistory of Nuʻu Refuge, Maui, Hawaiʻi. Marine Geology, 483, 107522. 10.1016/j.margeo.2025.107522

Frøslev, T. G., Kjøller, R., Bruun, H. H., Ejrnæs, R., Brunbjerg, A. K., Pietroni, C., & Hansen, A. J. (2017). Algorithm for post-clustering curation of DNA amplicon data yields reliable biodiversity estimates. Nature Communications, 8(1), 1188. 10.1038/s41467-017-01312-x

Fukami, T. (2015). Historical Contingency in Community Assembly: Integrating Niches, Species Pools, and Priority Effects. Annual Review of Ecology, Evolution, and Systematics, 46(Volume 46, 2015), 1–23. 10.1146/annurev-ecolsys-110411-160340

Galià-Camps, C., Baños, E., Pascual, M., Carreras, C., & Turon, X. (2023). Multidimensional variability of the microbiome of an invasive ascidian species. iScience, 26(10). 10.1016/j.isci.2023.107812

Gallardo, B., Clavero, M., Sánchez, M. I., & Vilà, M. (2016). Global ecological impacts of invasive species in aquatic ecosystems. Global Change Biology, 22(1), 151–163. 10.1111/gcb.13004

Garcés-Pastor, S., Coissac, E., Lavergne, S., Schwörer, C., Theurillat, J.-P., Heintzman, P. D., Wangensteen, O. S., Tinner, W., Rey, F., Heer, M., Rutzer, A., Walsh, K., Lammers, Y., Brown, A. G., Goslar, T., Rijal, D. P., Karger, D. N., Pellissier, L., Heiri, O., & Alsos, I. G. (2022). High resolution ancient sedimentary DNA shows that alpine plant diversity is associated with human land use and climate change. Nature Communications, 13(1), 6559. 10.1038/s41467-022-34010-4

Giguet-Covex, C., Pansu, J., Arnaud, F., Rey, P.-J., Griggo, C., Gielly, L., Domaizon, I., Coissac, E., David, F., Choler, P., Poulenard, J., & Taberlet, P. (2014). Long livestock farming history and human landscape shaping revealed by lake sediment DNA. Nature Communications, 5(1), 3211. 10.1038/ncomms4211

Goslar, T., Knaap, W. O. van der, Hicks, S., Andrič, M., Czernik, J., Goslar, E., Räsänen, S., & Hyötylä, H. (2005). Radiocarbon Dating of Modern Peat Profiles: Pre- and Post-Bomb 14C Variations in the Construction of Age-Depth Models. Radiocarbon, 47(1), 115–134. 10.1017/S0033822200052243

Guardiola, M., Uriz, M. J., Taberlet, P., Coissac, E., Wangensteen, O. S., & Turon, X. (2015). Deep-Sea, Deep-Sequencing: Metabarcoding Extracellular DNA from Sediments of Marine Canyons. PLOS ONE, 10(10), e0139633. 10.1371/journal.pone.0139633

Guillemaud, T., Ciosi, M., Lombaert, É., & Estoup, A. (2011). Biological invasions in agricultural settings: Insights from evolutionary biology and population genetics. Comptes Rendus Biologies, 334(3), 237–246. 10.1016/j.crvi.2010.12.008

Guo, Y., & Zhang, J. (2010). Numerical modelling of hydrodynamics in the Southampton Water. In Environmental Hydraulics. Volume 1. CRC Press.

Halpern, B. S., Frazier, M., Potapenko, J., Casey, K. S., Koenig, K., Longo, C., Lowndes, J. S., Rockwood, R. C., Selig, E. R., Selkoe, K. A., & Walbridge, S. (2015). Spatial and temporal changes in cumulative human impacts on the world’s ocean. Nature Communications, 6(1), 7615. 10.1038/ncomms8615

Haubrock, P. J., & Soto, I. (2023). Valuing the information hidden in true long-term data for invasion science. Biological Invasions, 25(8), 2385–2394. 10.1007/s10530-023-03091-7

Holman, L. E., Arfaoui, E. M. R., Pedersen, L. B., Farnsworth, W. R., Ascough, P., Butler, P., Guðmundsdóttir, E. R., Reynolds, D. J., Trofimova, T., Wilkin, J. T. R., Carøe, C., Frøslev, T. G., Harrison, R., Gopalakrishnan, S., Pedersen, M. W., Scourse, J., & Bohmann, K. (2025a). Ancient environmental DNA indicates limited human impact on marine biodiversity in pre-industrial Iceland. Philosophical Transactions B. 10.1098/rstb.2024.0031

Holman, L. E., Chng, Y., & Rius, M. (2022). How does eDNA decay affect metabarcoding experiments? Environmental DNA, 4(1), 108–116. 10.1002/edn3.201

Holman, L. E., Zampirolo, G., Gyllencreutz, R., Scourse, J., Frøslev, T., Carøe, C., Gopalakrishnan, S., Pedersen, M. W., & Bohmann, K. (2025b). Navigating Past Oceans: Comparing Metabarcoding and Metagenomics of Marine Ancient Sediment Environmental DNA. Molecular Ecology Resources, 25(6), e14086. 10.1111/1755-0998.14086

Hudson, J., Bourne, S. D., Seebens, H., Chapman, M. A., & Rius, M. (2022). The reconstruction of invasion histories with genomic data in light of differing levels of anthropogenic transport. Philosophical Transactions of the Royal Society B: Biological Sciences, 377(1846), 20210023. 10.1098/rstb.2021.0023

Jiang, X., Zhu, Z., Wu, J., Lian, E., Liu, D., Yang, S., & Zhang, R. (2022). Bacterial and Protistan Community Variation across the Changjiang Estuary to the Ocean with Multiple Environmental Gradients. Microorganisms, 10(5), 991. 10.3390/microorganisms10050991

Johnston, E. L., Dafforn, K. A., Clark, G. F., Rius, M., & Floerl, O. (2017). How Anthropogenic Activities Affect the Establishment and Spread of Non-Indigenous Species Post-Arrival. In S. J. Hawkins, A. J. Evans, A. C. Dale, L. B. Firth, D. J. Hughes, & I. P. Smith (Eds.), Oceanography and Marine Biology (1st ed., pp. 389–419). CRC Press. 10.1201/b21944-6

Katsanevakis, S., Wallentinus, I., Zenetos, A., Leppäkoski, E., Çinar, M. E., Oztürk, B., Grabowski, M., Golani, D., & Cardoso, A. C. (2014). Impacts of invasive alien marine species on ecosystem services and biodiversity: A pan-European review. Aquatic Invasions, 9(4), 391–423. 10.3391/ai.2014.9.4.01

Kennish, M. J. (2002). Environmental threats and environmental future of estuaries. Environmental Conservation, 29(1), 78–107. 10.1017/S0376892902000061

Kjær, K. H., Winther Pedersen, M., De Sanctis, B., De Cahsan, B., Korneliussen, T. S., Michelsen, C. S., Sand, K. K., Jelavić, S., Ruter, A. H., Schmidt, A. M. A., Kjeldsen, K. K., Tesakov, A. S., Snowball, I., Gosse, J. C., Alsos, I. G., Wang, Y., Dockter, C., Rasmussen, M., Jørgensen, M. E., … Willerslev, E. (2022). A 2-million-year-old ecosystem in Greenland uncovered by environmental DNA. Nature, 612(7939), 283–291. 10.1038/s41586-022-05453-y

Kumschick, S., & Richardson, D. M. (2013). Species-based risk assessments for biological invasions: Advances and challenges. Diversity and Distributions, 19(9), 1095–1105. 10.1111/ddi.12110

Kylander, M. E., Ampel, L., Wohlfarth, B., & Veres, D. (2011). High-resolution X-ray fluorescence core scanning analysis of Les Echets (France) sedimentary sequence: New insights from chemical proxies. Journal of Quaternary Science, 26(1), 109–117. 10.1002/jqs.1438

Lambert, G. (2007). Invasive sea squirts: A growing global problem. Journal of Experimental Marine Biology and Ecology, 342(1), 3–4. 10.1016/j.jembe.2006.10.009

Lanzén, A., Lekang, K., Jonassen, I., Thompson, E. M., & Troedsson, C. (2017). DNA extraction replicates improve diversity and compositional dissimilarity in metabarcoding of eukaryotes in marine sediments. PLOS ONE, 12(6), e0179443. 10.1371/journal.pone.0179443

Leigh, D. M., Hendry, A. P., Vázquez-Domínguez, E., & Friesen, V. L. (2019). Estimated six per cent loss of genetic variation in wild populations since the industrial revolution. Evolutionary Applications, 12(8), 1505–1512. 10.1111/eva.12810

Leray, M., Yang, J. Y., Meyer, C. P., Mills, S. C., Agudelo, N., Ranwez, V., Boehm, J. T., & Machida, R. J. (2013). A new versatile primer set targeting a short fragment of the mitochondrial COI region for metabarcoding metazoan diversity: Application for characterizing coral reef fish gut contents. Frontiers in Zoology, 10(1), 34. 10.1186/1742-9994-10-34

Levasseur, A. (2008). Observations and modelling of the variability of the Solent-Southampton Water estuarine system [Phd, University of Southampton]. https://eprints.soton.ac.uk/63761/

Levasseur, A., Shi, L., Wells, N. C., Purdie, D. A., & Kelly-Gerreyn, B. A. (2007). A three-dimensional hydrodynamic model of estuarine circulation with an application to Southampton Water, UK. Estuarine, Coastal and Shelf Science, 73(3), 753–767. 10.1016/j.ecss.2007.03.018

López-Legentil, S., Legentil, M. L., Erwin, P. M., & Turon, X. (2015). Harbor networks as introduction gateways: Contrasting distribution patterns of native and introduced ascidians. Biological Invasions, 17(6), 1623–1638. 10.1007/s10530-014-0821-z

Lotze, H. K., Lenihan, H. S., Bourque, B. J., Bradbury, R. H., Cooke, R. G., Kay, M. C., Kidwell, S. M., Kirby, M. X., Peterson, C. H., & Jackson, J. B. C. (2006). Depletion, Degradation, and Recovery Potential of Estuaries and Coastal Seas. Science, 312(5781), 1806–1809. 10.1126/science.1128035

Machida, R. J., Leray, M., Ho, S.-L., & Knowlton, N. (2017). Metazoan mitochondrial gene sequence reference datasets for taxonomic assignment of environmental samples. Scientific Data, 4(1), 170027. 10.1038/sdata.2017.27

Magurran, A. E., Deacon, A. E., Moyes, F., Shimadzu, H., Dornelas, M., Phillip, D. A. T., & Ramnarine, I. W. (2018). Divergent biodiversity change within ecosystems. Proceedings of the National Academy of Sciences, 115(8), 1843–1847. 10.1073/pnas.1712594115

Mahon, A. R., Jerde, C. L., Galaska, M., Bergner, J. L., Chadderton, W. L., Lodge, D. M., Hunter, M. E., & Nico, L. G. (2013). Validation of eDNA Surveillance Sensitivity for Detection of Asian Carps in Controlled and Field Experiments. PLOS ONE, 8(3), e58316. 10.1371/journal.pone.0058316

Martin, M. (2011). Cutadapt removes adapter sequences from high-throughput sequencing reads. EMBnet.Journal, 17(1), 10–12. 10.14806/ej.17.1.200

McMurdie, P. J., & Holmes, S. (2013). phyloseq: An R Package for Reproducible Interactive Analysis and Graphics of Microbiome Census Data. PLOS ONE, 8(4), e61217. 10.1371/journal.pone.0061217

Miura, O. (2007). Molecular genetic approaches to elucidate the ecological and evolutionary issues associated with biological invasions. Ecological Research, 22(6), 876–883. 10.1007/s11284-007-0389-5

Nelson, J. C. (2014). Species invasion in the marine fouling communities of British Columbia: Factors that influence invasion dynamics and how they may affect Botrylloides violaceus [University of British Columbia]. 10.14288/1.0167429

Nguyen, N.-L., Devendra, D., Szymańska, N., Greco, M., Angeles, I. B., Weiner, A. K. M., Ray, J. L., Cordier, T., De Schepper, S., Pawłowski, J., & Pawłowska, J. (2023). Sedimentary ancient DNA: A new paleogenomic tool for reconstructing the history of marine ecosystems. Frontiers in Marine Science, 10. 10.3389/fmars.2023.1185435

Nogué, S., Santos, A. M. C., Birks, H. J. B., Björck, S., Castilla-Beltrán, A., Connor, S., de Boer, E. J., de Nascimento, L., Felde, V. A., Fernández-Palacios, J. M., Froyd, C. A., Haberle, S. G., Hooghiemstra, H., Ljung, K., Norder, S. J., Peñuelas, J., Prebble, M., Stevenson, J., Whittaker, R. J., … Steinbauer, M. J. (2021). The human dimension of biodiversity changes on islands. Science, 372(6541), 488–491. 10.1126/science.abd6706

Ojaveer, H., Galil, B. S., Carlton, J. T., Alleway, H., Goulletquer, P., Lehtiniemi, M., Marchini, A., Miller, W., Occhipinti-Ambrogi, A., Peharda, M., Ruiz, G. M., Williams, S. L., & Zaiko, A. (2018). Historical baselines in marine bioinvasions: Implications for policy and management. PLoS ONE, 13(8), e0202383. 10.1371/journal.pone.0202383

Oksanen, J., Kindt, R., Legendre, P., Hara, B., Simpson, G., Solymos, P., Henry, M., Stevens, H., Maintainer, H., & Oksanen@oulu, jari. (2009). The vegan Package.

Osborne, K., & Poynton, H. (2019). Copper pollution enhances the competitive advantage of invasive ascidians. Management of Biological Invasions, 10(4), 641–656. 10.3391/mbi.2019.10.4.05

Pante, E., & Simon-Bouhet, B. (2013). marmap: A Package for Importing, Plotting and Analyzing Bathymetric and Topographic Data in R. PLOS ONE, 8(9), e73051. 10.1371/journal.pone.0073051

Parducci, L., Bennett, K. D., Ficetola, G. F., Alsos, I. G., Suyama, Y., Wood, J. R., & Pedersen, M. W. (2017). Ancient plant DNA in lake sediments. New Phytologist, 214(3), 924–942. 10.1111/nph.14470

Pawłowska, J., Lejzerowicz, F., Esling, P., Szczuciński, W., Zajączkowski, M., & Pawlowski, J. (2014). Ancient DNA sheds new light on the Svalbard foraminiferal fossil record of the last millennium. Geobiology, 12(4), 277–288. 10.1111/gbi.12087

Paxton, A. B., Runde, B. J., Smith, C. S., Lester, S. E., Vozzo, M. L., Saunders, M. I., Steward, D. N., Lemoine, H. R., Valdez, S. R., Gittman, R. K., Narayan, S., Allgeier, J., Morris, R. L., Nowacek, D. P., Seaman, W., Halpin, P. N., Angelini, C., & Silliman, B. R. (2025). Leveraging built marine structures to benefit and minimize impacts on natural habitats. BioScience, 75(2), 172–183. 10.1093/biosci/biae135

Pereira, H. M., Leadley, P. W., Proença, V., Alkemade, R., Scharlemann, J. P. W., Fernandez-Manjarrés, J. F., Araújo, M. B., Balvanera, P., Biggs, R., Cheung, W. W. L., Chini, L., Cooper, H. D., Gilman, E. L., Guénette, S., Hurtt, G. C., Huntington, H. P., Mace, G. M., Oberdorff, T., Revenga, C., … Walpole, M. (2010). Scenarios for Global Biodiversity in the 21st Century. Science, 330(6010), 1496–1501. 10.1126/science.1196624

Pérez, V., Liu, Y., Wong, W. W., Kessler, A., Cook, P. L. M., Zawadzki, A., Moore, N. E., Kurte, L., Child, D., Hotchkis, M., Weyrich, L. S., & Lintern, A. (2023). Using sedimentary prokaryotic communities to assess historical changes in the Gippsland Lakes. Freshwater Biology, 68(11), 1839–1858. 10.1111/fwb.14182

Pineda, M. C., McQuaid, C. D., Turon, X., López-Legentil, S., Ordóñez, V., & Rius, M. (2012). Tough Adults, Frail Babies: An Analysis of Stress Sensitivity across Early Life-History Stages of Widely Introduced Marine Invertebrates. PLOS ONE, 7(10), e46672. 10.1371/journal.pone.0046672

Platin, R., & Shenkar, N. (2023). Can stand the heat – ecology of the potentially invasive ascidian *Styela plicata* along the Mediterranean coast of Israel. Frontiers in Marine Science, 10. 10.3389/fmars.2023.1159231

Ratnasingham, S., & Hebert, P. D. N. (2013). A DNA-Based Registry for All Animal Species: The Barcode Index Number (BIN) System. PLOS ONE, 8(7), e66213. 10.1371/journal.pone.0066213

Reitzel, A. M., Darling, J. A., Sullivan, J. C., & Finnerty, J. R. (2008). Global population genetic structure of the starlet anemone Nematostella vectensis: Multiple introductions and implications for conservation policy. Biological Invasions, 10(8), 1197–1213. 10.1007/s10530-007-9196-8

Ricciardi, A., Blackburn, T. M., Carlton, J. T., Dick, J. T. A., Hulme, P. E., Iacarella, J. C., Jeschke, J. M., Liebhold, A. M., Lockwood, J. L., MacIsaac, H. J., Pyšek, P., Richardson, D. M., Ruiz, G. M., Simberloff, D., Sutherland, W. J., Wardle, D. A., & Aldridge, D. C. (2017). Invasion Science: A Horizon Scan of Emerging Challenges and Opportunities. Trends in Ecology & Evolution, 32(6), 464–474. 10.1016/j.tree.2017.03.007

Rius, M., & Turon, X. (2020). Phylogeography and the Description of Geographic Patterns in Invasion Genomics. Frontiers in Ecology and Evolution, 8. 10.3389/fevo.2020.595711

Rius, M., Turon, X., Bernardi, G., Volckaert, F. A. M., & Viard, F. (2015). Marine invasion genetics: From spatio-temporal patterns to evolutionary outcomes. Biological Invasions, 17(3), 869–885. 10.1007/s10530-014-0792-0

Sakata, M. K., Yamamoto, S., Gotoh, R. O., Miya, M., Yamanaka, H., & Minamoto, T. (2020). Sedimentary eDNA provides different information on timescale and fish species composition compared with aqueous eDNA. Environmental DNA, 2(4), 505–518. 10.1002/edn3.75

Sardain, A., Sardain, E., & Leung, B. (2019). Global forecasts of shipping traffic and biological invasions to 2050. Nature Sustainability, 2(4), 274–282. 10.1038/s41893-019-0245-y

Schnell, I. B., Bohmann, K., & Gilbert, M. T. P. (2015). Tag jumps illuminated – reducing sequence-to-sample misidentifications in metabarcoding studies. Molecular Ecology Resources, 15(6), 1289–1303. 10.1111/1755-0998.12402

Seebens, H., Blackburn, T. M., Dyer, E. E., Genovesi, P., Hulme, P. E., Jeschke, J. M., Pagad, S., Pyšek, P., Winter, M., Arianoutsou, M., Bacher, S., Blasius, B., Brundu, G., Capinha, C., Celesti-Grapow, L., Dawson, W., Dullinger, S., Fuentes, N., Jäger, H., … Essl, F. (2017). No saturation in the accumulation of alien species worldwide. Nature Communications, 8(1), 14435. 10.1038/ncomms14435

Sharifi, A. R., Croudace, I. W., & Austin, R. L. (1991). Benthic foraminiferids as pollution indicators in Southampton Water, southern England, U.K. Journal of Micropalaeontology, 10(1), 109–113. 10.1144/jm.10.1.109

Simberloff, D., Martin, J.-L., Genovesi, P., Maris, V., Wardle, D. A., Aronson, J., Courchamp, F., Galil, B., García-Berthou, E., Pascal, M., Pyšek, P., Sousa, R., Tabacchi, E., & Vilà, M. (2013). Impacts of biological invasions: What’s what and the way forward. Trends in Ecology & Evolution, 28(1), 58–66. 10.1016/j.tree.2012.07.013

Sorte, C. J. B., Williams, S. L., & Carlton, J. T. (2010). Marine range shifts and species introductions: Comparative spread rates and community impacts. Global Ecology and Biogeography, 19(3), 303–316. 10.1111/j.1466-8238.2009.00519.x

Stager, J. C., Sporn, L. A., Johnson, M., & Regalado, S. (2015). Of Paleo-Genes and Perch: What if an “Alien” Is Actually a Native? PLOS ONE, 10(3), e0119071. 10.1371/journal.pone.0119071

Stat, M., Huggett, M. J., Bernasconi, R., DiBattista, J. D., Berry, T. E., Newman, S. J., Harvey, E. S., & Bunce, M. (2017). Ecosystem biomonitoring with eDNA: Metabarcoding across the tree of life in a tropical marine environment. Scientific Reports, 7(1), 12240. 10.1038/s41598-017-12501-5

Stuble, K. L., & Souza, L. (2016). Priority effects: Natives, but not exotics, pay to arrive late. Journal of Ecology, 104(4), 987–993. 10.1111/1365-2745.12583

Taberlet, P., Bonin, A., Zinger, L., & Coissac, E. (2018). Environmental DNA: For Biodiversity Research and Monitoring. Oxford University Press.

Tang, C. Q., Leasi, F., Obertegger, U., Kieneke, A., Barraclough, T. G., & Fontaneto, D. (2012). The widely used small subunit 18S rDNA molecule greatly underestimates true diversity in biodiversity surveys of the meiofauna. Proceedings of the National Academy of Sciences, 109(40), 16208–16212. 10.1073/pnas.1209160109

Tavener, L. E. (1950). The Port of Southampton. Economic Geography. https://www.tandfonline.com/doi/abs/10.2307/141262

Torres, A., Rodríguez-Cabal, M. A., & Núñez, M. A. (2022). Do not come late to the party: Initial success of nonnative species is contingent on timing of arrival of co-occurring nonnatives. Biological Invasions, 24(2), 557–573. 10.1007/s10530-021-02660-y

Turbelin, A. J., Diagne, C., Hudgins, E. J., Moodley, D., Kourantidou, M., Novoa, A., Haubrock, P. J., Bernery, C., Gozlan, R. E., Francis, R. A., & Courchamp, F. (2022). Introduction pathways of economically costly invasive alien species. Biological Invasions, 24(7), 2061–2079. 10.1007/s10530-022-02796-5

Turner, A. (2010). Marine pollution from antifouling paint particles. Marine Pollution Bulletin, 60(2), 159–171. 10.1016/j.marpolbul.2009.12.004

van Leeuwen, J. F. N., Froyd, C. A., van der Knaap, W. O., Coffey, E. E., Tye, A., & Willis, K. J. (2008). Fossil Pollen as a Guide to Conservation in the Galápagos. Science, 322(5905), 1206–1206. 10.1126/science.1163454

Vilà, M., & Hulme, P. E. (2017). Non-native Species, Ecosystem Services, and Human Well-Being. In M. Vilà & P. E. Hulme (Eds.), Impact of Biological Invasions on Ecosystem Services (pp. 1–14). Springer International Publishing. 10.1007/978-3-319-45121-3_1

Wagstaff, M. (2017). Life history variation of an invasive species Botrylloides violaceus (Oka, 1927) between novel coastal habitats in the Gulf of Maine. Aquatic Invasions, 12(1), 43–51. 10.3391/ai.2017.12.1.05

Walentowitz, A., Lenzner, B., Essl, F., Strandberg, N., Castilla-Beltrán, A., Fernández-Palacios, J. M., Björck, S., Connor, S., Haberle, S. G., Ljung, K., Prebble, M., Wilmshurst, J. M., Froyd, C. A., de Boer, E. J., de Nascimento, L., Edwards, M. E., Stevenson, J., Beierkuhnlein, C., Steinbauer, M. J., & Nogué, S. (2023). Long-term trajectories of non-native vegetation on islands globally. Ecology Letters, 26(5), 729–741. 10.1111/ele.14196

Wang, Y., Pedersen, M. W., Alsos, I. G., De Sanctis, B., Racimo, F., Prohaska, A., Coissac, E., Owens, H. L., Merkel, M. K. F., Fernandez-Guerra, A., Rouillard, A., Lammers, Y., Alberti, A., Denoeud, F., Money, D., Ruter, A. H., McColl, H., Larsen, N. K., Cherezova, A. A., … Willerslev, E. (2021). Late Quaternary dynamics of Arctic biota from ancient environmental genomics. Nature, 600(7887), 86–92. 10.1038/s41586-021-04016-x

Wangensteen, O. S., Palacín, C., Guardiola, M., & Turon, X. (2018). DNA metabarcoding of littoral hard-bottom communities: High diversity and database gaps revealed by two molecular markers. PeerJ, 6, e4705. 10.7717/peerj.4705

Wauchope, H. S., Amano, T., Geldmann, J., Johnston, A., Simmons, B. I., Sutherland, W. J., & Jones, J. P. G. (2021). Evaluating Impact Using Time-Series Data. Trends in Ecology & Evolution, 36(3), 196–205. 10.1016/j.tree.2020.11.001

Weigand, H., Beermann, A. J., Čiampor, F., Costa, F. O., Csabai, Z., Duarte, S., Geiger, M. F., Grabowski, M., Rimet, F., Rulik, B., Strand, M., Szucsich, N., Weigand, A. M., Willassen, E., Wyler, S. A., Bouchez, A., Borja, A., Čiamporová-Zaťovičová, Z., Ferreira, S., … Ekrem, T. (2019). DNA barcode reference libraries for the monitoring of aquatic biota in Europe: Gap-analysis and recommendations for future work. Science of The Total Environment, 678, 499–524. 10.1016/j.scitotenv.2019.04.247

Wickham, H. (2011). Ggplot2. WIREs Computational Statistics, 3(2), 180–185. 10.1002/wics.147

Yebra, D. M., Kiil, S., & Dam-Johansen, K. (2004). Antifouling technology—Past, present and future steps towards efficient and environmentally friendly antifouling coatings. Progress in Organic Coatings, 50(2), 75–104. 10.1016/j.porgcoat.2003.06.001

Ytreberg, E., Bighiu, M. A., Lundgren, L., & Eklund, B. (2016). XRF measurements of tin, copper and zinc in antifouling paints coated on leisure boats. Environmental Pollution, 213, 594–599. 10.1016/j.envpol.2016.03.029

Zarcero, J., Antich, A., Rius, M., Wangensteen, O. S., & Turon, X. (2024). A new sampling device for metabarcoding surveillance of port communities and detection of non-indigenous species. iScience, 27(1), 108588. 10.1016/j.isci.2023.108588

Zinger, L., Bonin, A., Alsos, I. G., Bálint, M., Bik, H., Boyer, F., Chariton, A. A., Creer, S., Coissac, E., Deagle, B. E., De Barba, M., Dickie, I. A., Dumbrell, A. J., Ficetola, G. F., Fierer, N., Fumagalli, L., Gilbert, M. T. P., Jarman, S., Jumpponen, A., … Taberlet, P. (2019). DNA metabarcoding—Need for robust experimental designs to draw sound ecological conclusions. Molecular Ecology, 28(8), 1857–1862. 10.1111/mec.15060

